# Redistribution of ancestral functions underlies the evolution of venom production in marine predatory snails

**DOI:** 10.1101/2024.09.09.612013

**Authors:** Giulia Zancolli, Maria Vittoria Modica, Nicolas Puillandre, Yuri Kantor, Agneesh Barua, Giulia Campli, Marc Robinson-Rechavi

## Abstract

Venom-secreting glands are highly specialised organs evolved throughout the entire animal kingdom to synthetise and secrete toxins for predation and defence. Venom is extensively studied for its toxin components and application potential; yet, how animals become venomous remains poorly understood. Venom systems therefore offer a unique opportunity to understand the molecular mechanisms underlying functional innovation. Here, we conducted a multi-species multi-tissue comparative transcriptomics analysis of 12 marine predatory gastropods, including species with venom glands and species with homologous non-venom producing glands, to examine how specialised functions evolve through gene expression changes. We found that while the venom gland specialised for the mass production of toxins, its homologous glands retained the ancestral digestive functions. The functional divergence and specialisation of the venom gland was achieved through a redistribution of its ancestral digestive functions to other organs, specifically the oesophagus. This entailed concerted expression changes and accelerated transcriptome evolution across the entire digestive system. The increase in venom gland secretory capacity was achieved through the modulation of an ancient secretory machinery, particularly genes involved in endoplasmic reticulum stress and unfolded protein response. This study shifts the focus from the well-explored evolution of toxins to the lesser-known evolution of the organ and mechanisms responsible for venom production. As such, it contributes to elucidating the molecular mechanisms underlying organ evolution at a fine evolutionary scale, highlighting the specific events that lead to functional divergence.

## Introduction

Across diverse branches of the animal kingdom, organisms have independently evolved the ability to produce and deliver venom, a cocktail of bioactive toxin molecules. In many venomous animals, these toxins are synthetised in specialised exocrine organs known as venom glands (Casewell et al. 2013; Schendel et al. 2019). While significant research has focused on the molecular evolution of toxins and venom composition, the molecular mechanisms underlying the evolution of venom-producing organs and their specialised function remains poorly understood (Zancolli and Casewell 2020). Recent genomics studies in snakes, for instance, have highlighted how regulatory networks are co-opted to drive high toxin gene expression in venom glands, notably through the involvement of trans-regulatory factors from the extracellular signal-regulated kinase (ERK) and unfolded protein response (UPR) pathways (Perry et al. 2020; Perry et al. 2022; Westfall et al. 2023). Moreover, the upregulation of UPR and endoplasmic reticulum (ER) stress pathways in the venom glands of several distinct venomous taxa suggests that similar molecular solutions may have convergently evolved across venomous lineages to support the high secretory demands of toxin production (Perry et al. 2020; Barua and Mikheyev 2021; Zancolli et al. 2022). However, a broader understanding of how venom glands become highly specialised and optimised for the efficient mass production of toxins is still lacking, particularly outside of snake models.

Investigating homologous structures with divergent functions offers unique insights into the genetic basis of functional innovation, as demonstrated in large-scale evolutionary studies on vertebrate limbs (Zuniga 2015) and feathers (Benton et al. 2019). Among non-vertebrates, the mid-oesophageal glands of marine predatory snails in the subclass Caenogastropoda (Fig. 1) provide an excellent system to examine, at a finer evolutionary scale, the specific events that led to the emergence of new physiological functions. In some caenogastropods, like those in the family Naticidae, the oesophageal gland consist of a dilated section of the oesophageal wall involved in secreting digestive enzymes and mucus (Reid and Friesen 1980; Lobo-da-Cunha 2019). By contrast, in many caenogastropods of the order Neogastropoda, this glandular section of the oesophagus has evolved into a distinct organ known as the gland of Leiblein, which connects to the oesophagus via a duct (Modica and Holford 2010; Lobo-da-Cunha 2019). Ultrastructural observations in species from the families Muricidae and a Nassariidae suggest that the gland of Leiblein plays a role in food processing, particularly nutrient absorption and storage (Andrews and Thorogood 2005). In the superfamily Conoidea, which comprises 18 families (Abdelkrim et al. 2018) including the well-known cone snails (family Conidae) (Nguyen et al. 2023), the oesophageal gland has undergone further modifications into a long, convoluted duct that secretes a mixture of hundreds of primarily neurotoxic peptides, mainly known as ‘conotoxins’ (Modica and Holford 2010; Abdelkrim et al. 2018; Morales Duque et al. 2019). The venom gland is attached to a large muscular bulb that contracts to push venom through the duct and into the buccal cavity, where a modified radula injects venom into prey or predators (Modica and Holford 2010; Safavi-Hemami et al. 2010; Salisbury et al. 2010). Interestingly, some neogastropods, such as those in the family Mitridae (Harasewych 2009) and Terebridae (Holford et al. 2009), have entirely lost the mid-oesophageal gland.

**Figure 1:**
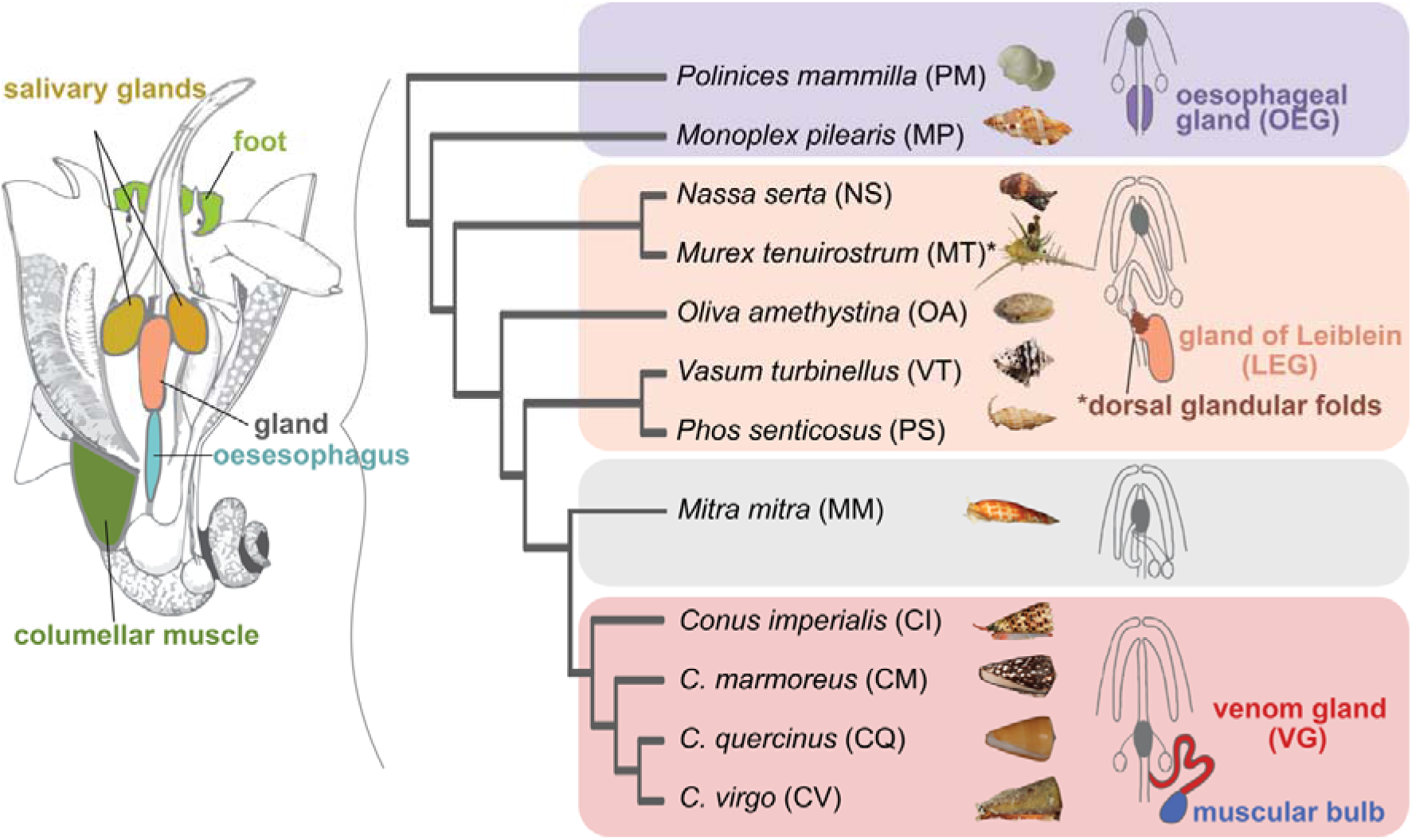
Species and tissues investigated. Overview of the anatomy of a marine snail with the color-coded sampled tissues and the phylogeny of the species used in this study based on Fedosov et al. (2024). The abbreviations used for the tissues and species used in other figures are shown. The anatomy drawing is modified after (Modica and Holford 2010), while the schematic of the foregut apparatus of caenogastropods with their mid-oesophageal glands was modified after (Page 2011).

The evolution of venom production from digestive-related functions in these snails represents a remarkable example of functional specialisation, shifting the gland’ s secretion targets from internal digestive roles (e.g., lysosomal activity) (Andrews and Thorogood 2005) to external roles that affect other organisms (receptor binding in prey) (Morales Duque et al. 2019). This transition likely provided significant adaptive advantages, enabling cone snails to diversify their diet to include fast-moving organisms, such as fish, while also offering a defence against powerful predators (Olivera et al. 2012). However, the processes through which an organ originally dedicated to digestive functions transformed into a specialised toxin-producing factory remain unclear.

Here, we analyse gene expression data from the mid-oesophageal glands and other tissues of 12 marine caenogastropod species to investigate the link between transcriptome evolution and functional divergence. By studying the genetic underpinnings of venom gland evolution, we aim to uncover how specific genes, pathways, and regulatory networks drive organ specialisation to meet the distinct physiological demands of venom production. Our study addresses the following key questions: (i) Do mid-oesophageal glands across species share similar gene expression profiles, given their common origin? (ii) Considering the venom gland’s unique function, does its transcriptome evolve more rapidly compared to the other mid-oesophageal glands? (iii) Which gene expression changes led to the evolution of toxin production in the mid-oesophageal gland?

To answer these questions, we first conducted species-level analyses to characterise sets of overexpressed genes and to delineate the functional specialisation of each gland type. We then performed between-species comparisons to explore the gene expression dynamics that led to the evolution of venom production. Our findings reveal that the venom gland has a markedly distinct gene expression profile compared to its homologous organs, which are more similar to each other. Genes encoding secreted proteins in venom glands are expressed at exceptionally high levels, far exceeding those of other tissues. This specialisation for toxin secretion was achieved through modulation of a conserved secretory machinery, while ancestral digestive functions were redistributed to other organs. This shift involved high evolutionary rates not only in the venom gland itself but across the entire digestive system, suggesting concerted changes that underscore the adaptive flexibility of these organisms.

## Results

### Summary of species and tissues investigated

We sampled ten species of Neogastropoda and two outgroup species within the same subclass Caenogastropoda (Fig. 1). The two outgroup species, the Cymatidae *Monoplex pilearis* (Linnaeus, 1758) and the Naticidae *Polinices mammilla* (Linnaeus, 1758), possess a simple oesophageal gland (OEG) attached to the oesophagus, representing the ancestral state. Among the Neogastropoda, two species from the Muricidae family, *Nassa serta* (Bruguière, 1789) and *Murex tenuirostrum* (Lamarck, 1822), have a gland of Leiblein (LEG), with *Murex tenuirostrum* also possessing dorsal glandular folds on the mid-oesophagus, often referred to as glande framboisée (Lobo Cunha 2019). Additionally, we sampled the gland of Leiblein in the Olividae *Oliva amethystina* (Rőding, 1798), the Vasidae *Vasum turbinellus* (Linnaeus, 1758), and the Nassaridae *Phos senticosus* (Linnaeus, 1758). The venomous species, possessing a venom gland (VG), belonged to the family Conidae, including *Conus imperialis* (Linnaeus, 1758), *C. marmoreus* (Linnaeus, 1758), *C. virgo* (Linnaeus, 1758), and *C. quercinus* (Lightfoot, 1786). We also collected tissue samples from the Mitridae *Mitra mitra* (Linnaeus, 1758), which lacks a mid-oesophageal gland (Harasewych 2009). Besides the glands, we sampled either the foot or the columellar muscle, oesophagus, salivary glands, dorsal glandular folds in *M. tenuirostrum*, and muscular venom bulb in cone snails.

### Sequencing and de novo assembly statistics

A total of 150 libraries were sequenced, yielding an average of 45 million reads per library across 12 species, with an average of three biological replicates per species (supplementary dataset S1). We generated *de novo* assemblies by pooling all libraries within each species and processing them through our quality-filtering pipeline (see Methods). After filtering, the assemblies contained between 26,215 and 59,431 annotated, non-redundant protein-coding genes, with an average of 40,192 sequences per assembly (supplementary dataset S2). The completeness of these assemblies ranged from 82% to 93%, with an average of 89%. Prior to data analysis, we evaluated the quality of read alignment. Samples with low mapping rates or those that did not cluster with other samples of the same tissue type in a Principal Component Analysis (PCA) plot were excluded (see Methods). After this quality filtering step, we retained a total of 140 samples (supplementary fig. S1, Supplementary Material online).

### Within-species analysis

#### Characterisation of tissue-specific gene sets

To better understand the functional specialisation of the mid-oesophageal glands, we identified sets of genes that were overexpressed in each organ. Genes were classified as tissue-specific if their expression in a tissue was at least twice that of the second most highly expressed tissue. On average, 14% (range: 10-19%) of genes were tissue-specific, with 43% (range: 38-51%) of these genes exclusively expressed in one tissue. Tissue specificity was consistent across organs and species, although we observed a higher number of tissue-specific genes in the venom glands compared to the glands of Leiblein and oesophageal glands (Fig. 2a). The smallest set observed was the oesophagus of *M. tenurostrum* (N = 318). Notably, this species possesses dorsal glandular folds on the oesophagus (Fig. 1). The tissue-specificity of the gene sets was further validated by differential expression analysis using a likelihood ratio test across all tissues (supplementary text 1, Supplementary Material online).

**Figure 2:**
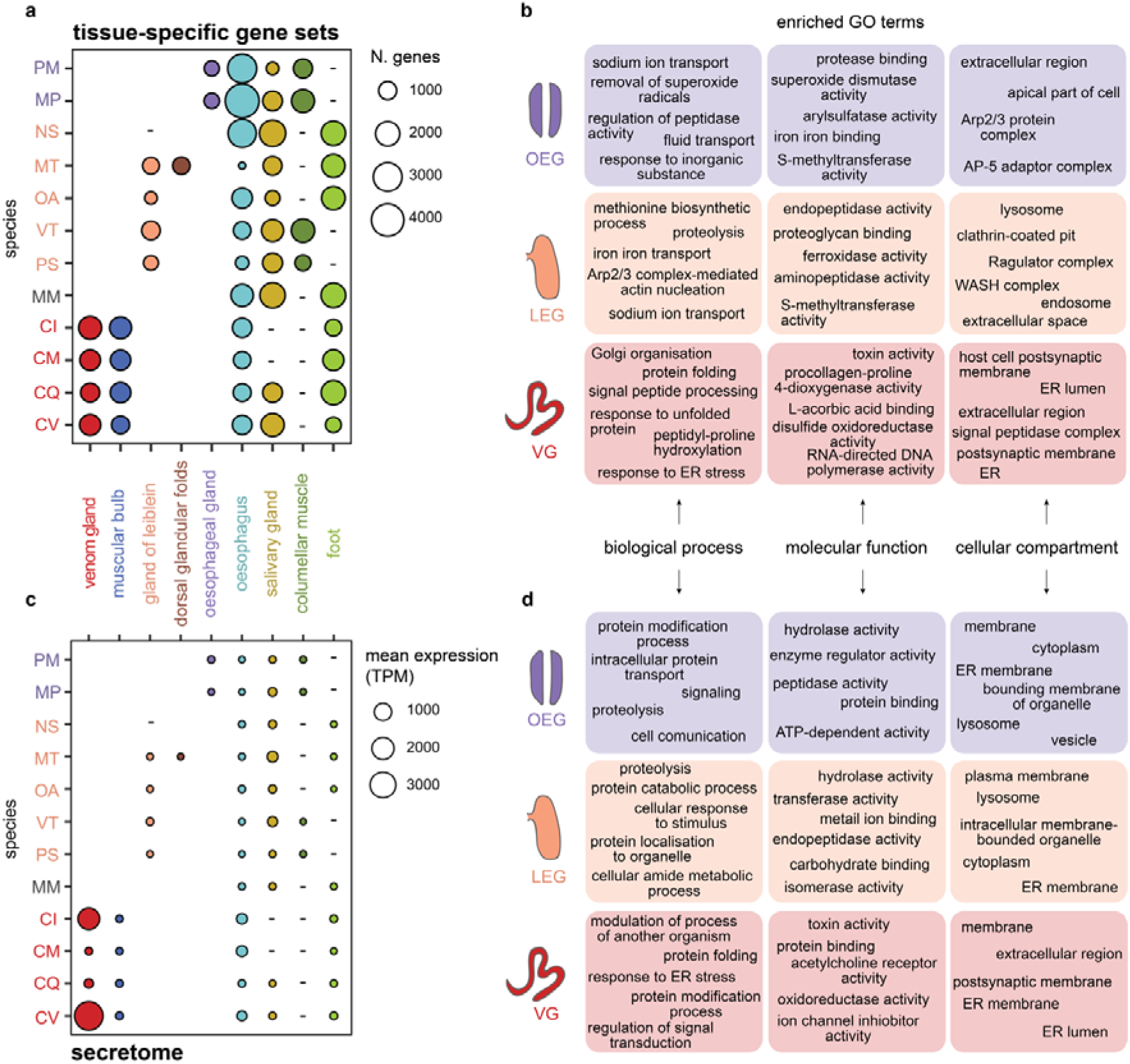
Overview of tissue-specific gene sets and gland secretomes. a) Number of genes within each tissue-specific set across all organs (x-axis) and species (y-axis). Species abbreviations as in Fig. 1. The venom glands have a significant higher number of tissue-specific genes (mean N = 1,424) compared to the gland of Leiblein (mean N = 771; t = 3.6, df = 6, p = 0.005) and oesophageal gland (mean N = 699; t = 5.6, df =3, p = 0.005). Missing tissue samples are marked with a “-”. b) Gene Ontology (GO) enrichment results of the OEG-, LEG-, and VG-specific gene sets. c) Mean expression levels as Transcript per Million (TPM) of genes possessing a signal peptide, therefore comprising the secretomes, expressed in the organs (x-axis) of each species (y-axis). Species abbreviations as in Fig. 1. Missing tissue samples are marked with a “-”. d) GO enrichment results of the oesophageal gland, gland of Leiblein, and venom gland secretomes.

Gene Ontology (GO) enrichment analysis of the tissue-specific gene sets revealed marked differences between the mid-oesophageal glands (Fig. 2b), as well as lineage-specific patterns (supplementary text 2 and figs. S2-S10, Supplementary Material online). In OEG-specific gene sets, we observed enrichment for terms related to intracellular trafficking and communication (e.g., ‘AP-5 adaptor complex’) and transmembrane transport. Additionally, terms related to detoxification, homeostasis, and response to external stimuli (e.g., ‘response to inorganic substance’, ‘superoxide dismutase activity’) were enriched (supplementary figs. S2-S4, Supplementary Material online). In contrast, gene sets specific to the gland of Leiblein across multiple species were enriched for terms related to intracellular digestion, particularly protein digestion and absorption, with several lysosome-related terms (e.g., ‘endopeptidase activity’, ‘WASH complex’), aligning with previous ultrastructural studies (Andrews and Thorogood 2005). Terms related to iron homeostasis and metabolism were also enriched. However, *O. amethystina* showed some unique enrichments, including extracellular rather than intracellular compartment terms, reflecting possible lineage-specific adaptations linked to diet (supplementary figs. S5-S7, Supplementary Material online). In venom glands, GO terms associated with toxin and neurotoxic activity (e.g., ‘host cell postsynaptic membrane’) were over-represented in all species, as expected (Fig. 2b). Additionally, we found terms related to protein synthesis (e.g., ‘signal peptide processing’, ‘Golgi organisation’) and to post-translational modifications (e.g., ‘hydroxylation’, ‘disulfide oxidoreductase activity’). Notably, terms related to ER stress and UPR were also enriched (e.g., ‘response to unfolded proteins’, ‘response to ER stress’) (supplementary figs. S8-S10, Supplementary Material online).

In summary, while the non-venomous mid-oesophageal glands are primarily involved in digestion, homeostasis, absorption, and storage, the homologous venom gland has specialised into a factory for toxin synthesis and secretion. Despite these differences, all three gland types shared enriched terms related to iron homeostasis and metabolism (e.g., ‘iron binding’) and methylation (‘methionine adenosyltransferase activity’, ‘betaine-homocysteine S-methyltransferase activity’). Additionally, we found terms related to hormone response across all three gland types, including ‘thyroid hormone generation’ in OEG- and LEG-specific gene sets, and ‘response to thyroglobulin triiodothyronine’ in VG-specific sets.

#### Characterisation of the glands’ secretomes

Given the fundamental role of secretion in the evolution of the venom gland, we analysed the ‘secretome’ by assessing the diversity and expression levels of genes predicted to have a signal peptide with SignalP (Teufel et al. 2022).

The number of expressed genes encoding secreted proteins was relatively consistent across tissue types, although it was higher in the venom gland (mean N = 565.25) compared to the gland of Leiblein (mean N = 354.25) and the oesophageal gland (mean N = 379.5) (supplementary fig. S11a, Supplementary Material online). When focussing on tissue-specific genes, the venom gland secretome was also more diverse than the other glands (supplementary fig. S11b, Supplementary Material online). Interestingly, the salivary glands did not show particularly high diversity despite being exocrine glands like the venom gland (supplementary fig. S11, Supplementary Material online).

In terms of expression levels (mean TPM), the differences between tissues were more pronounced (Fig. 2c). Gene expression was generally higher in venom glands, although it varied among cone snail species (Fig. 2c). This variation could reflect genuine lineage-specific differences, technical factors, or differences in the venom replenishment circle at the time of sampling. However, the latter is unlikely, as individuals were sampled randomly and thus would not all be at the same point in their replenishment cycle. Additionally, all specimens were kept for 1-3 days in captivity prior dissection to minimise environmental variation. We also did not observe ingested prey in any stomachs, suggesting that the last feeding – and thus venom expulsion - did not occur close to the time of dissection for any individual. In non-venomous species, the highest expression levels were observed in the salivary glands. Interestingly, in the oesophagus, lower expression levels were observed in non-venomous species (mean = 21 TPM) compared to venomous species (mean = 260; t = - 3, df = 3, p = 0.02), suggesting higher secretory activity in the latter.

As anticipated, the secretomes of the oesophageal gland and gland of Leiblein were enriched in hydrolases and peptidases, enzymes essential for digestive processes (Fig. 2d). Additionally, *O. amethystina* showed enrichment in ‘toxin activity’ (see next section), while *M. pilearis* was enriched in terms related to communication and transport (e.g., ‘metal ion transport’). Although the salivary glands also expressed hydrolases and peptidases, these enzymes were extracellular, whereas those in the gland of Leiblein were primarily intracellular, consistent with enrichment in cellular compartments like ‘lysosome’ and ‘organelle lumen”. In contrast, the venom gland secretomes were dominated by toxins released in the extracellular space and of genes involved in the ER function (Fig. 2d).

#### Identification and characterisation of conotoxins

Cone snail venom is composed primarily of small, disulfide-rich peptides known as conotoxins. The number of putative conotoxin transcripts predicted in the venomous species’ assemblies was consistent with previous findings for cone snail venom gland *de novo* transcriptomes (Abalde et al. 2018; Dutt et al. 2019), with 150-250 toxins predicted per species. Notably, conotoxin-like sequences were also identified in other species, with numbers ranging from 33 in *M. mitra* to 88 in *O*. *amethystina*. However, when restricting the analysis to sequences with both a signal peptide and a predicted conotoxin domain, the numbers were greatly reduced, ranging from 16 to 108 in venomous species, and 7 to 28 in non-venomous species.

As expected, conotoxins were predominantly expressed in the venom gland (supplementary fig. S12, Supplementary Material online). A few were also highly and specifically expressed in the salivary glands, consistent with previous reports in other cone snail species (Fedosov et al. 2023). In non-venomous species, predicted conotoxins were expressed at much lower levels and across multiple tissues, although a trend towards expression in salivary glands was observed (supplementary figs. S13-S14, Supplementary Material online).

It is important to note that the prediction tool used, ConoPrec (Kaas et al. 2012), is design to predict only conoidean toxins. Consequently, toxins found outside Conoidea, like echotoxin (Shiomi et al. 2002), were not predicted, leading to an underestimation of the toxic potential of gastropod salivary glands (Ponte and Modica 2017). Our focus on conotoxins stems from their critical role as the primary weapon of cone snails, whose massive gene expansion and diversification likely drove the evolution of the venom gland.

### Between-species analysis

#### Orthogroup assignment

We assigned 330,491 genes (68% of the total) to 41,720 orthogroups (OGs), with 2,588 OGs shared across all species. For comparative transcriptomics, we created a multi-species expression matrix using the 2,588 OGs. Since many OGs contained multiple genes per species, we selected a representative gene for each OG using two methods: 1) randomly selecting a single transcript’s TPM value, and 2) calculating the mean TPM across all the transcripts within an OG. Both approaches produced similar outcomes, therefore the results presented here are based on the first method. Results from the second approach are provided in the Supplementary Material online.

#### Transcriptome similarity and shared tissue specificity between organs and species

To determine whether homologous glands share similar global gene expression profiles or if their functional specialisations align them more closely with non-homologous organs, we analysed whole transcriptome similarity patterns by means of correlation matrix and PCA. Overall, samples primarily clustered by tissue type (Fig. 3a, supplementary figs. S15-S16, Supplementary Material online). The oesophageal glands and glands of Leiblein grouped together, while the venom glands clustered with the salivary glands rather than with the homologous counterparts. Interesting patterns were observed when comparing tissue specificity across species. We found that the oesophageal glands on average shared more tissue-specific OGs with the glands of Leiblein (mean N = 61, sd = 17.7) and the oesophagus of the glandless *M. mitra* (mean N. = 56, sd = 10.6) rather than with each other (mean N. = 41, sd = 0). This suggests substantial variation between the two OEG-species, as corroborated by the GO enrichment analysis (supplementary figs. S2-S4, Supplementary Material online). The glands of Leiblein, in contrast, shared more tissue-specific OGs among themselves (mean N. 103, sd = 45.9), and with the oesophagus of *M. mitra* (mean N. = 100, sd = 38.2). In contrast, venom glands shared more tissue-specific OGs among themselves (mean N = 111, sd = 20.4) and with the salivary glands of *M. mitra* (mean N = 71, 11.5), while very few with the oesophageal glands (mean N = 11, sd = 3.6) and glands of Leiblein (mean N = 16, 7.7). In summary, while the oesophageal gland and gland of Leiblein share more similar transcriptomes, the venom gland markedly diverged from its homologous organs and converged towards the other exocrine organ, the salivary glands

**Figure 3.**
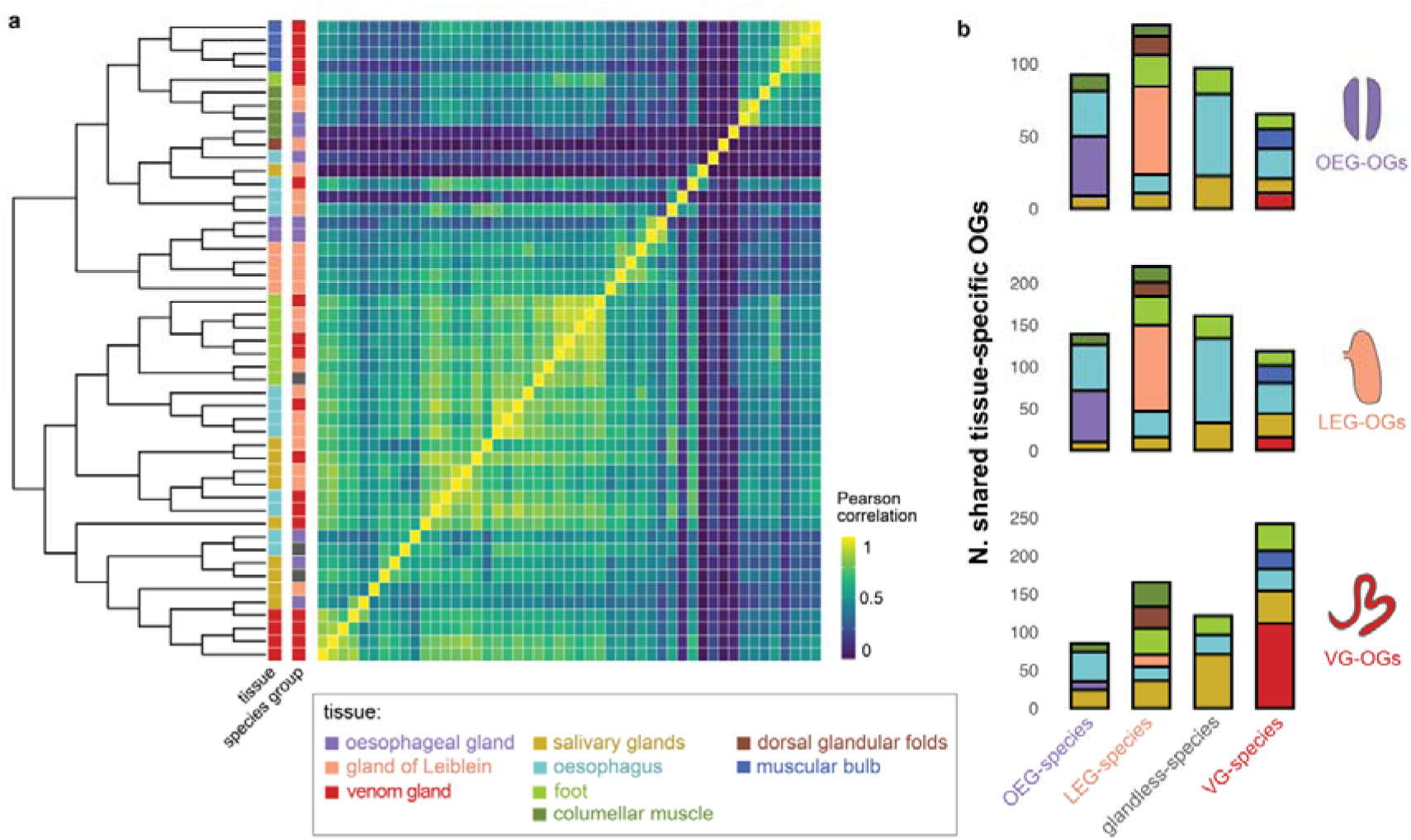
Transcriptome similarity and shared tissue specificity. a) Heatmap of Pearson correlation coefficients between tissues and species. The expression tree was made using neighbour-joining based on the correlation matrix. The samples are colored based on the tissue and the gland type that the species possess. b) Average number of OEG-, LEG-, and VG-specific OGs shared with other tissue-specific OG sets, where species have been grouped by their gland type (OEG-, LEG-, VG-species and glandless).

For subsequent analyses, we defined OGs as tissue-specific if they were upregulated in the same tissue of at least two species, resulting in 41 OEG-specific OGs, 343 LEG-specific OGs, and 405 VG-specific OGs.

#### Rates of gene expression evolution

Given the marked functional divergence of the venom gland, we hypothesised that its transcriptome evolves faster than that of its homologous glands. We tested this hypothesis using CAGEE (Bertram et al. 2023), which employs a bounded Brownian motion model to estimate the most likely value of the evolutionary rate parameter ( ^2^) consistent with an ultrametric species tree and observed expression values at the tip of the tree. We ran CAGEE for the mid-oesophageal glands, salivary glands, and oesophagus, as these tissues were sampled across most species representing all three gland types.

We evaluated four different evolutionary models (supplementary fig. S17, Supplementary Material online). The first model estimated a single rate ^2^. The second model estimated two distinct rates, one for the venomous clade and one for all other species. In the third model, species were grouped by gland type and the rates were estimated for each group separately. The final model also calculated three rates but assigned them randomly across the phylogeny. Overall, the third model had the best fit (Table 1, supplementary table S1, Supplementary Material online). Notably, the model with the poorest fit was the first one, indicating that some degree of variation in evolutionary rates, even if random, is more consistent with the data than assuming a uniform rate across all lineages. When comparing across lineages, the venom gland showed the highest ^2^ value, supporting our hypothesis of accelerated evolution in the venom gland relative to the other homologous glands. A similar trend was observed in other organs, with higher evolutionary rates in venomous species than non-venomous ones. However, when comparing across organs, none of the glands had the highest ^2^ value, suggesting that other organs also underwent accelerated evolution, even more so than the mid-oesophageal glands, aligning with the hypothesis of concerted evolution across the entire digestive system.

**Table 1.**
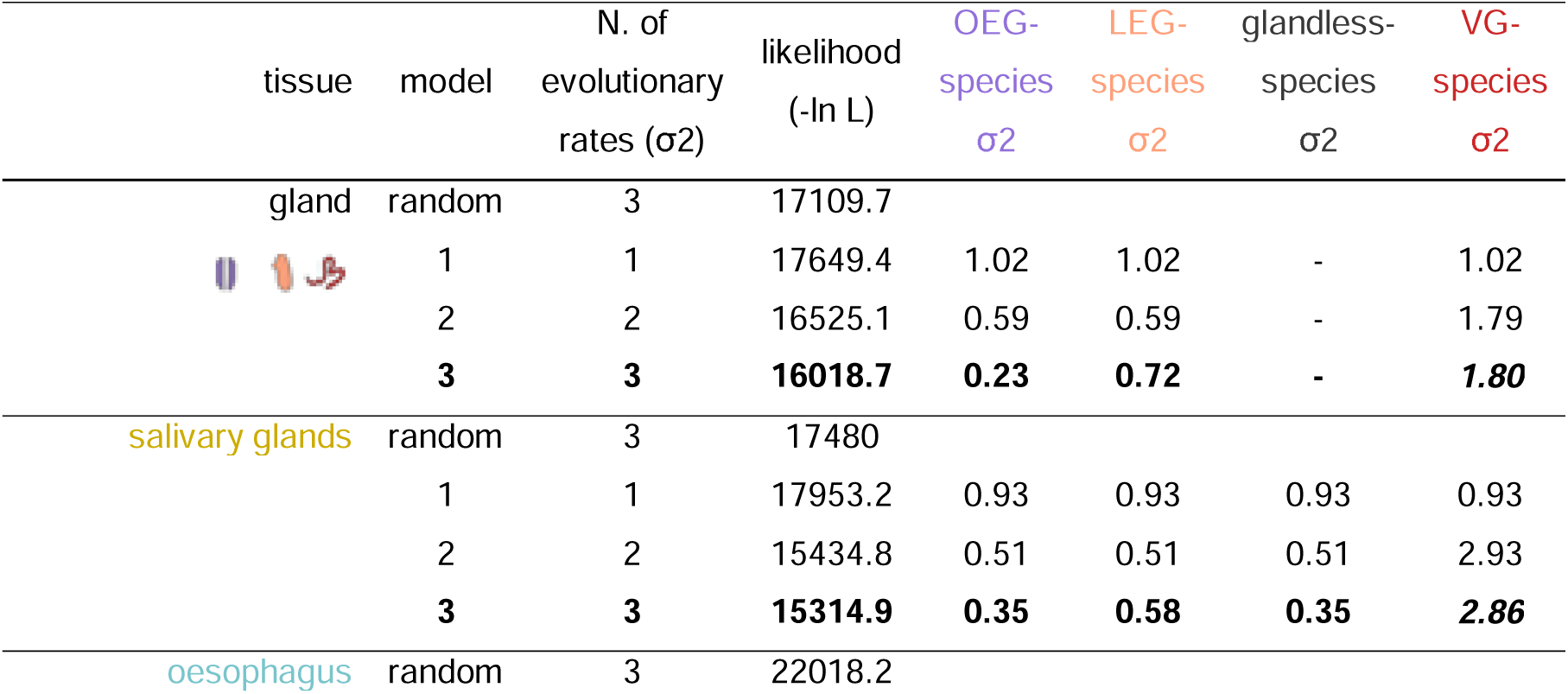
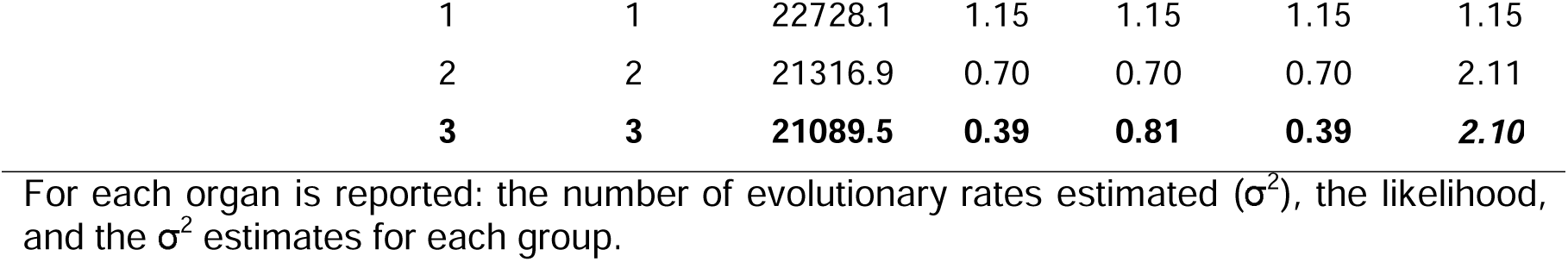
Evolutionary rates for the tested models.

For each organ is reported: the number of evolutionary rates estimated (σ^2^), the likelihood, and the σ^2^ estimates for each group.

#### Expression changes of tissue-specific orthogroups along the phylogeny

Given the better fit of the third evolutionary model, we used this one for ancestral state reconstruction to assess the number and direction of expression changes at each node of the phylogenetic tree (Fig. 4b). We observed substantial changes at node 18, which leads to the glandless *M. mitra* and the venomous clade (Fig. 4a, supplementary fig. S18, Supplementary Material online). At this node, many OEG- and LEG-specific OGs showed a marked decrease in expression in the ancestral gland, coupled with an increase in expression in the oesophagus and salivary glands. Conversely, VG-specific OGs underwent upregulation in the gland as well as across other tissues (Fig. 4a).

**Figure 4.**
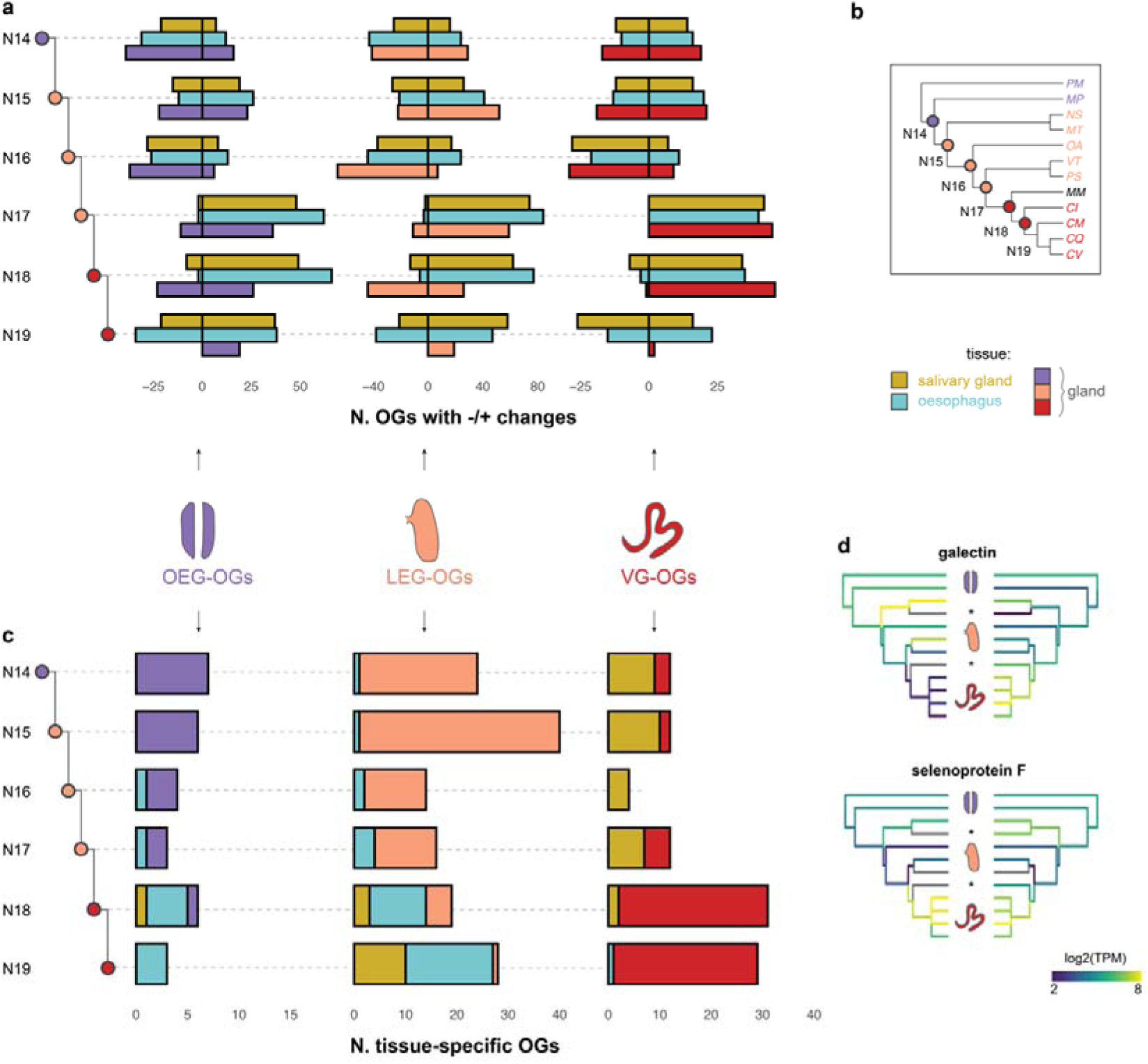
Gene expression dynamics across the phylogeny. a) Number of OEG-, LEG, and VG-specific OGs decreasing and increasing their expression levels in the ancestral salivary glands, oesophagus, or mid-oesophageal gland at each internal node of the gastropod phylogeny. The expression changes were calculated based on gene expression reconstruction at each node of the phylogeny. b) Species phylogeny with the number of the internal nodes. c) Number of OEG-, LEG-, and VG-specific OGs which were tissue-specific in the ancestral mid-oesophageal gland, salivary glands, and oesophagus at each node of the phylogeny. d) Ancestral reconstruction of gene expression of a galectin (OG2631) and selenoprotein F (OG465) in the mid-oesophageal gland and oesophagus. The missing tissues are marked with a *.

Among the OEG- and LEG-specific OGs, the strongest downregulation in the gland was a galectin, which simultaneously showed the highest upregulation in the oesophagus (Fig. 4d), alongside with a member of the ependymin family. The top three VG-specific OGs that were upregulated in the gland included an integral membrane protein of the DAD family, a disulfide-isomerase, and a selenoprotein F (Fig. 4d), all involved in the ER.

We then utilised the ancestral state reconstruction from CAGEE to calculate tissue specificity at each node, akin to our approach for extant species. The goal was to understand whether gland-specific OGs were tissue-specific in the ancestral lineages. Our findings reveal that some OEG- and LEG-specific OGs were gland-specific in ancestral lineages until node 18, where they shifted mainly to the oesophagus (Fig. 4c, supplementary fig. S18, Supplementary Material online). The galectin and ependymin mentioned earlier were among the genes that transitioned to oesophagus specificity, alongside other genes involved in gluconeogenesis. In contrast, VG-specific OGs were primarily salivary gland-specific in early nodes but shifted to VG-specific at node 18 (Fig. 4c). These shifts suggest that the organs of the digestive system had to adapt to the new function, or loss of, the mid-oesophageal gland.

## Discussion

Understanding how organs evolve is central to studying functional innovation. Our investigation into marine gastropod mid-oesophageal glands provides a unique perspective on the specific events driving the emergence of specialised functions like venom production. Venom glands in animals are highly specialised exocrine organs dedicated primarily to toxin synthesis and secretion (Schendel et al. 2019). Consistent with this, our functional enrichment analyses reveals that venom glands in cone snails are distinguished by exceptionally highly upregulation of protein secretion pathways, mirroring findings in other lineages (Perry et al. 2020; Barua and Mikheyev 2021; Zancolli et al. 2022). In contrast, genes specifically expressed in the gland of Leiblein, and, to a lesser extent, the oesophageal gland, are associated with intracellular digestion, particularly of proteins, as well as uptake and storage of lipids and carbohydrates (Andrews and Thorogood 2005). The oesophageal gland, in particular, also appears to play roles in homeostasis, transport of fluids and nutrients, and secretion of digestive enzymes (Reid and Friesen 1980).

The phylogenetic distribution of these mid-oesophageal glands, combined with the functions inferred from their transcriptomes suggest a scenario of concerted functional and morphological evolution closely linked to their trophic ecology. Initially, the role of these glands in nutrient absorption and intracellular digestion would have required prey to be either pre-digested by salivary enzymes or naturally soft-tissues. For example, species in the Muricidae family that drill holes into prey and consume soft tissues have a well-developed gland of Leiblein (Andrews and Thorogood 2005), and their salivary glands open near the tip of the proboscis, enabling immediate interaction of salivary products with prey (Kantor 1996). As gastropods diversified their diets and some species adopted macrophagous feeding strategies, the glands evolved accordingly, trending towards reduction or even loss. For instance, Buccinoidea species ingest prey whole or in large chunks (Kantor 2003) and possess a simple gland structure (Andrews and Thorogood 2005). In cases where the gland no longer functioned in absorption, it either disappeared, as in the Mitridae, or specialised for new functions, as seen in conoideans like cone snails. While it is often proposed that venom evolved to enable animals to capture and consume larger prey, our findings suggest an intriguing reversal in marine gastropods: an initial shift towards macrophagus feeding may have paved the way for venom evolution rather than vice versa.

The functional transformation of the mid-oesophageal gland in Conoidea was facilitated by coordinated adaptations in other digestive organs, which shifted their functions accordingly. Our ancestral state reconstruction reveals extensive changes in gene expression across the phylogeny, reflecting the dynamic nature of this system and the adaptation of these snails to diverse ecological niches and feeding strategies. Notably, the venomous clade experienced more substantial expression changes and higher evolutionary rates than non-venomous species, consistent with the drastic functional divergence of the venom gland from its homologous counterparts. However, the venom gland itself did not exhibit higher evolutionary rates than the homologous glands, nor higher rates compared to other organs. This suggests concerted evolution across the digestive systems, with other organs co-evolving to support the gland’s new function. In the ancestor of the venomous species and the glandless *M. mitra* (node 18), several genes became downregulated in the gland while upregulated in the oesophagus (Fig. 4a, c). Among these were genes involved in gluconeogenesis, such as galectin and PEPCK, indicating that the ancestral gland’s role in producing energy from non-carbohydrate substrates was reassigned to other digestive tissues. This concerted adaptation is particularly evident in the glandless species, where many OEG- and LEG-specific genes are now oesophagus-specific (Fig. 3b), especially those related to lysosomal functions. Mitridae snails feed on soft-bodied Sipuncula worms which they may ingest whole (Taylor 1989; Taylor 1993), or by pumping the worm’s viscera (West 1990), including coelomic fluids and eggs (West 1991), into the buccal cavity. In such cases, nutrients may be pre-digested by salivary enzymes, allowing the oesophagus to take over functions previously performed by the gland of Leiblein.

During conoidean evolution, the mid-oesophageal gland lost its original digestive functions but gained enhanced secretory capacity, primarily through the modulation of genes involved in pre-existing secretory pathways. Our analysis identified key upregulated genes in the ancestral venom gland (node 18) that encode proteins active in the ER, including DAD1, which is critical for N-glycosylation and protein translocation (Zhang et al. 2016), disulfide isomerases, and a selenoprotein F likely involved in ER protein folding quality control (Takeda et al. 2014). Additionally, genes associated with the UPR and ER stress response were enriched in cone snail venom glands, echoing patterns seen across other venomous lineages (Perry et al. 2020; Zancolli et al. 2022). This finding is significant for two reasons: first, gastropod venom glands reaffirm the trend of convergent transcriptomic evolution in venom glands across Metazoa (Zancolli et al. 2022). Second, the upregulation of these pathways appears unique to venom glands, as neither the extant homologous glands not the ancestral organ show this pattern based on transcriptome reconstruction. The strong upregulation of the UPR pathway in venom glands is notable. Even when compared to other exocrine organs with high secretory demands, such as salivary glands (this study) or the pancreas (Perry et al. 2020), venom glands show much higher expression levels. This indicates that heightened UPR expression is a distinct adaptation that evolved specifically within venom glands to support their unique physiological demands and directly controlling toxin expression. Studies in snakes have led to a model where venom production activates the UPR, creating a positive feedback loop that enhances venom production through up-regulation and binding of UPR transcription factors targeting toxin genes (Perry et al. 2020; Perry et al. 2022; Westfall et al. 2023).

In earlier nodes of the tree, OEG- and LEG-specific genes were expressed in the ancestral gland, while VG-specific genes were found in salivary glands - a pattern still evident in modern species (Fig. 3, Fig. 4c). This convergence between venom and salivary glands is logical, as both are exocrine organs secreting products into the extracellular environment. Notably, the salivary glands of some neogastropods secrete toxins to immobilise prey (Ponte and Modica 2017), thereby expanding their function. In these cases, salivary secretions serve both endogenous roles (e.g., pre-digestion enzymes) and exogenous functions (e.g., toxins), indicating a functional convergence between these two non-homologous organs. Interestingly, the salivary glands of the glandless species *M. mitra* share several tissue-specific genes with the venom glands. Unlike other snails, *Mitra* has salivary ducts that open at the tip of the epiproboscis, an extendible muscular rod within the proboscis (Harasewych 2009), likely facilitating the direct delivery of secretions to prey (West 1990). We observed overexpression of cysteine-rich secreted proteins, peptidases, and serine proteases in *Mitra*’s salivary glands, which suggests roles in tissue degradation and potential toxin activity. Despite these functional similarities, salivary glands retained their original digestive role and only secondarily adopted a toxin-secreting function, while the venom gland became fully specialised solely for the latter. A plausible explanation for why the salivary glands did not evolve into specialised venom glands lies, again, in the diet – Mitridae snails primarily feed on Sipuncula worms, a relatively inactive and scarcely targeted prey group. This reduced competition may have lessen the selective pressure to develop a specialised venom apparatus, unlike cone snails that faced greater competition and adapted to prey on fast-moving organisms that required paralysing toxins for successful capture.

## Conclusions

This study provides new insights into the genetic basis of functional innovation by examining the mid-oesophageal glands of marine gastropods, with a particular focus on the evolution of venom production. Our results indicate that while mid-oesophageal glands share a common origin, they have diverged in gene expression profiles and functions, shaped by adaptations to different feeding strategies. In cone snails, ancestral digestive functions of the mid-oesophageal gland were relocated to other digestive tissues in a process of concerted evolution, enabling the venom gland to specialise exclusively in toxin production through modulation of pathways related to secretion and cellular stress management. Overall, this study underscores the link between transcriptome evolution and functional divergence and identify the specific events leading to the emergence of new physiological functions.

## Materials and Methods

### Sample collection and sequencing

Between two and five individuals from Neogastropoda, and two outgroup species, were collected in Koumac, New Caledonia, under permit N°609011-55 /2019/DEPART/JJC. Individuals were kept for a minimum of one day and a maximum of three days in aquaria with fresh sea water before dissection. Salivary glands, foot, columellar muscle, oesophagus, oesophageal gland, gland of Leiblein, dorsal glandular folds (only in *M. tenuirostrum*), venom gland, and muscular bulb (only in cone snails) were dissected and preserved in RNA*later* (Invitrogen) (supplementary dataset S1).

Tissue samples were homogenised with Trizol (Invitrogen) and total RNA purified with the PureLink^TM^ RNA Mini kit (Invitrogen) with an additional DNase I treatment following manufacturer’s protocol. cDNA libraries were constructed using the NEBnext Ultra II Directional RNA Library Preparation Kit with polyA selection and dUTP method (New England BioLabs) and sequenced on an Illumina NovaSeq 2×150bp. Raw paired-end reads were checked with FastQC 0.11.9 (Andrews 2010), quality-filtered and trimmed with FastP 0.20.1 (Chen et al. 2018).

### De novo assembly and annotation

All the reads from a species were pooled to generate *de novo* transcriptome assemblies using rnaSpades 3.15.2 (Bushmanova et al. 2019). To reduce assembly size and redundancy, and remove spurious transcripts, we adopted a series of filtering steps. First, we translated the transcripts to amino acid sequences using Borf 1.2.1 (Signal and Kahlke 2021) and kept only those with a complete open read frame. Second, we compared the translated sequences with BlastP (Altschul et al. 1997) against a suite of databases (downloaded on 30.07.2021) including UniprotKB/Swiss-Prot (The UniProt Consortium 2023), a set of 11 Gastropoda genomes (supplementary table S2, Supplementary Material online), Conoserver (Kaas et al. 2012) and Tox-Prot (Jungo et al. 2012). Protein domains were identified with PSIblast (Altschul et al. 1997) against the Pfam (Mistry et al. 2021) and Cdd (Wang et al. 2022) databases. Only hits with evalue < 1e-05 were retained. Additionally, we annotated signal peptides with signalP 6.0 (Teufel et al. 2022). Putative conotoxins were predicted with the ConoPrec tool available on ConoServer (Kaas et al. 2012). All sequences with a UniprotKB/Swiss-Prot hit to a non-Metazoa organism were removed. We then trimmed the retained transcripts to their coding region, reduced redundancy by clustering sequences with 98% or more identical nucleotide sequences with CD-HIT-EST 4.6 (Li and Godzik 2006), and kept only transcripts with more than one read count in at least one library. Finally, we evaluated assembly completeness using Omark on the webserver omark.omabrowser.org (Nevers et al. 2024).

For each species we performed GO annotation by combining the annotation from Pannzer2 (Törönen et al. 2018) and DeepGOPlus 1.0.2 (Kulmanov and Hoehndorf 2020). Based on the score values distribution, we used 0.3 as a score threshold for both methods.

### Within-species analysis

#### Expression levels

We mapped all libraries to the respective species assembly with Kallisto 0.48.0 (Bray et al. 2016) with 100 bootstrap and quantified gene expression using the package Sleuth 0.30.1 (Pimentel et al. 2017) using R 4.2.2 (R Core Team 2019). We employed a suite of quality control steps to identify and remove outliers. First, we analysed read count distribution with *vioplot* 0.4.0 and removed samples with particularly different distributions, generally with read counts lower than the average. Then, we utilise dimensionality reduction techniques, including PCA (*dudi_PCA*) and multidimensional scaling (*plotMDS*) to verify that samples were clustering by tissue type. If a sample had unusual clustering, i.e., it was far from the others of the same tissue, we excluded it. Normalised estimated counts and TPM abundances were then re-calculated after outlier libraries’ removal. As in some species samples still tended to cluster by individual rather than tissue type, we account for the specimen effect using an empirical Bayes method implemented via the *ComBat_seq* function in the package sva 3.46.0 (Leek et al. 2012) which specifically targets RNA-Seq data. We build a full model as ∼ specimen + tissue_type, and a reduced model as ∼ specimen.

#### Tissue-specific gene sets

For each species, we identified tissue-specific genes based on their fold change (FC) calculated as the ratio between the TPM value of the first most-highly expressed tissue and the TPM value of the second-most highly expressed tissue. A gene with TPM ≥ 2 and FC ≥ 2 was classified as specific of the top tissue. We validated our FC method by confirming that the tissue-specific genes identified were also significantly differentially expressed when analysed using the likelihood ratio test in Sleuth. Differences in the number of tissue-specific genes between glands were tested by means of one-tailed t-tests. Functional enrichment of tissue-specific gene sets was performed with TopGO 2.50.0 (Alexa and Rahnenfuhrer 2019) using the *elim* algorithm and Fisher test. The foreground was the list of tissue-specific genes while the background included all the genes expressed in that species.

### Between-species analysis

#### Orthogroup expression matrix

Amino acid sequences were assigned to orthogroups (OGs) with the OrthoDB standalone pipeline OrthoLoger 3.0.2 (Kuznetsov et al. 2023) using default parameters. Since most OGs included more than one gene per species (i.e., one-to-many or many-to-many orthologs), we randomly selected one representative sequence for each OG in each species and used the TPM value estimated for that gene as the orthogroup expression value. This method was shown to be robust (Zancolli et al. 2022). Alternatively, we calculated the mean TPM values across all genes belonging to the same OG. All samples were then merged into a multi-species multi-tissue matrix and the expression levels corrected for the species effect using *ComBat_seq*. All the downstream analyses reported in the main text are based on the random matrix, while the results from the mean-based matrix are reported in Supplementary Material online.

To have an overview of global transcriptome similarity across tissues and species, we calculated pairwise distances as 1-Spearman correlation and used it to reconstruct a gene expression tree using the neighbour-joining method. Additionally, we calculated the proportion of shared OGs among tissue-specific genes. For downstream analyses, OGs were classified tissue-specific if at least one gene within that OG was specific in a tissue, and if it was found specific in at least two species.

#### Gene expression evolution analysis

We analysed changes in gene expression with the program CAGEE 1.1.1 (Bertram et al. 2023) which uses Brownian motion to model gene expression across a phylogenetic tree. The tree was derived from the phylogeny of Fedosov et al. (2024) by excluding the families not encompassed in this study and retaining only the branches most closely related to the species that we examined. The rates of expression changes (σ^2^) were calculated for the mid-oesophageal glands, salivary glands, and oesophagus separately. We fit a series of nested models in which σ^2^ varies across branches of the species tree to test for different hypotheses as outlined in the results. Ancestral transcriptomes at inner nodes reconstructed in the best fit model were used to calculate ancestral tissue specificity for each OG as we did for extant species.

#### Evolution of novel genes

To test whether new genes evolved along with the venom gland, we examined the number of OGs that were found exclusively in the venomous species, as well as whether venom gland-specific OGs had, on average, a higher number of genes in venomous species compared to the other species (i.e., mean OG size in VG-species > mean OG size of all the other species groups).

## Acknowledgments

This work has received funding from the European Union’s Horizon 2020 research and innovation programme under Marie Skłodowska-Curie grant agreement 845674 to G.Z, and grant agreement 865101 to N.P. We are grateful to Philippe Buchet, the team of the expeditions Koumac 2.1 and 2.3 of the Muséum National d’Histoire Naturelle, Paris, and the Province Nord and Government of New Caledonia for providing the logistic support and expertise, David Massemin and Patrick Marti for their skills in finding the species needed. We thank Alexander Fedesov for providing the phylogenetic tree, Consolée Aletti for support with the RNA extraction, the personnel of the Genomic Technologies Facility and Genewiz for the RNA-seq experiment, and members of the Robinson Rechavi’s lab for bioinformatics support and scientific discussions. The first version of the manuscript was greatly improved thanks to the feedback of two anonymous reviewers and additional reviewers through Qeios.

## Author contributions

G.Z., M.V.M., N.P., Y.K., and M.R.-R. conceived the study design; Y.K., and M.V.M perform the dissections; G.Z. generated the RNA-seq data, perfomed data analysis, and wrote the manuscript; A.B. performed the GO annotation and G.C. the orthogroup assignment. All the authors contributed to the interpretation and discussion of the results, and to the final version of the manuscript.

## Competing interests

The authors declaire no competing interests.

## Data availability

The RNA-seq data generated in this study have been deposited in the NCBI SRA archive with the accession number PRJNA1158673. Additional data generated in this study are available at 10.5281/zenodo.13685166. Additional information are provided as Supplementary files.

## Notes

### Competing Interest Statement

The authors have declared no competing interest.

### Summary of Updates

We have improved the overall flow and clarity of the text, added a conclusion section, and removed the part on novel genes and transposable elements.

