## Supplementari Materials online for "Redistribution of ancestral functions underlies the evolution of venom production in marine predatory snails"

### Table of contents

**Supplementary Text 1**

- Differential expression analysis

**Supplementary Text 2**

- Functional enrichment of tissue-specific gene sets

**Supplementary References**

**Supplementary Figures**

- Supplementary Figures S1 to S19

**Supplementary Tables**

- Supplementary Tables S1 to S2

**Legends for Dataset S1 and S2**

### Supplementary Text 1

##### Differential expression analysis

The specificity of the gene sets identified with the fold change approach was confirmed by differential expression (DE) analysis with a likelihood ratio test across all tissues. Briefly, the likelihood ration test was computed with the command *sleuth_lrt* with a reduced model, and a full model which fits the tissue type while accounting for the specimen effect. On average, 8.5% (5-12%) of all the expressed genes were differentially expressed across tissues. However, these identified gene sets did not include genes which were exclusively expressed in one tissue (i.e., private genes) as the pipeline in the Sleuth package automatically removes genes with TPM zero across all samples but in one tissue type. Furthermore, DE analysis included genes that were upregulated in more than one tissue. Therefore, all the genes that are highly specific to only one tissue, and which therefore most relevant for functional divergence, were excluded by the DE method. Between 48-75% of the DE genes identified with the likelihood ratio test were identified with the fold change (FC) method. The DE genes which were not listed as FC genes corresponded to genes which were upregulated in more than one tissue type.

### Supplementary Text 2

##### Functional enrichment of tissue-specific gene sets

*Mid-oesophageal glands*

In the oesophageal glands we found Gene Ontology (GO) terms related mainly to transport, for instance transmembrane transport, Arp2/3 protein complex, apical part of cell, and AP-5 adaptor complex. Terms related to breaking down of macromolecules were also enriched, such as arylsulfatase activity and carboxypeptidase activity. There were differences between the two species. For instance, in *P. mammilla* we found terms related to endosomes such as WASH complex and endosome membrane, as well as lysosomes (clathrin coat, BLOC-1 complex), and glutathione transferase which is important for detoxification.

In the glands of Leiblein we found GO terms related to intracellular digestion, especially of proteins, and absorption including proteolysis, collagen catabolic process, endopeptidase activity, lysosome, clathrin-coated pit and Ragulator complex.

In venom glands, over-represented GO terms were related to protein synthesis such as signal peptide processing, Golgi organization, peptidyl-proline hydroxilation, disulfide oxidoreductase activity. Of course, there were many terms related to toxins such as modulation of process of another organism, synaptic transmission, acetylcholin receptor activity, postsynaptic membrane. The endoplasmic reticulum stress and unfolded protein response pathways, which are typically upregulated in venom gland transcriptomes, were also enriched.

*Salivary glands*

Regarding other important organs, salivary glands in species with oesophageal glands had enrichment of terms similar to those of the venom gland, such as protein targeting to endoplasmic reticulum and signal peptide processing.

In species with the gland of Leiblein, the enriched GO terms were much more variables, with not that many terms in common across all species. Nonetheless, we found an overrepresentation of terms related to defense response to bacteria and transcription initiation, Golgi apparatus, endopeptidase and oxidoreductase activity.

Salivary glands of the venomous species were enriched in terms related to transport, especially fluids (sodium ion, phosphate ion, water channel activity), and secretion of enzymes to break down macromolecules such as carbonate dehydratase activity and lipase activity. These terms summarise the classic functions of salivary glands.

In the glandless species, enriched terms were related to translation and ribosomes, signal peptide processing, protein targeting ER, aminopeptidase activity, glycosylation, and disulfide reductase activity.

*Dorsal glandular folds of Murex tenuirostrum*

The dorsal glandular folds of *M. tenuirostrum* were enriched with terms related to ribosomes and translation (e.g., 'regulation of translation', 'rRNA processing', 'RNA binding'), specifically the initial stages of ribosomal biogenesis and assembly. Additional terms included endosome, ER, signaling, protein maturation, splicing and Golgi apparatus suggesting secretion, but also terms related to immune response.

*Muscular venom bulb*

The muscular bulb, which is part of the venom apparatus of cone snails, was enriched with of terms related to myosin, collagen, cytoskeleton, and calcium binding, confirming its muscular nature. Additionally, we found arginine kinase activity, as it was previously found in proteomics investigation (Safavi-Hemami et al. 2010). This enzyme has been reported in muscles undergoing burst contraction, as it requires high levels of ATP generated via arginine kinase activity (Safavi-Hemami et al. 2010).

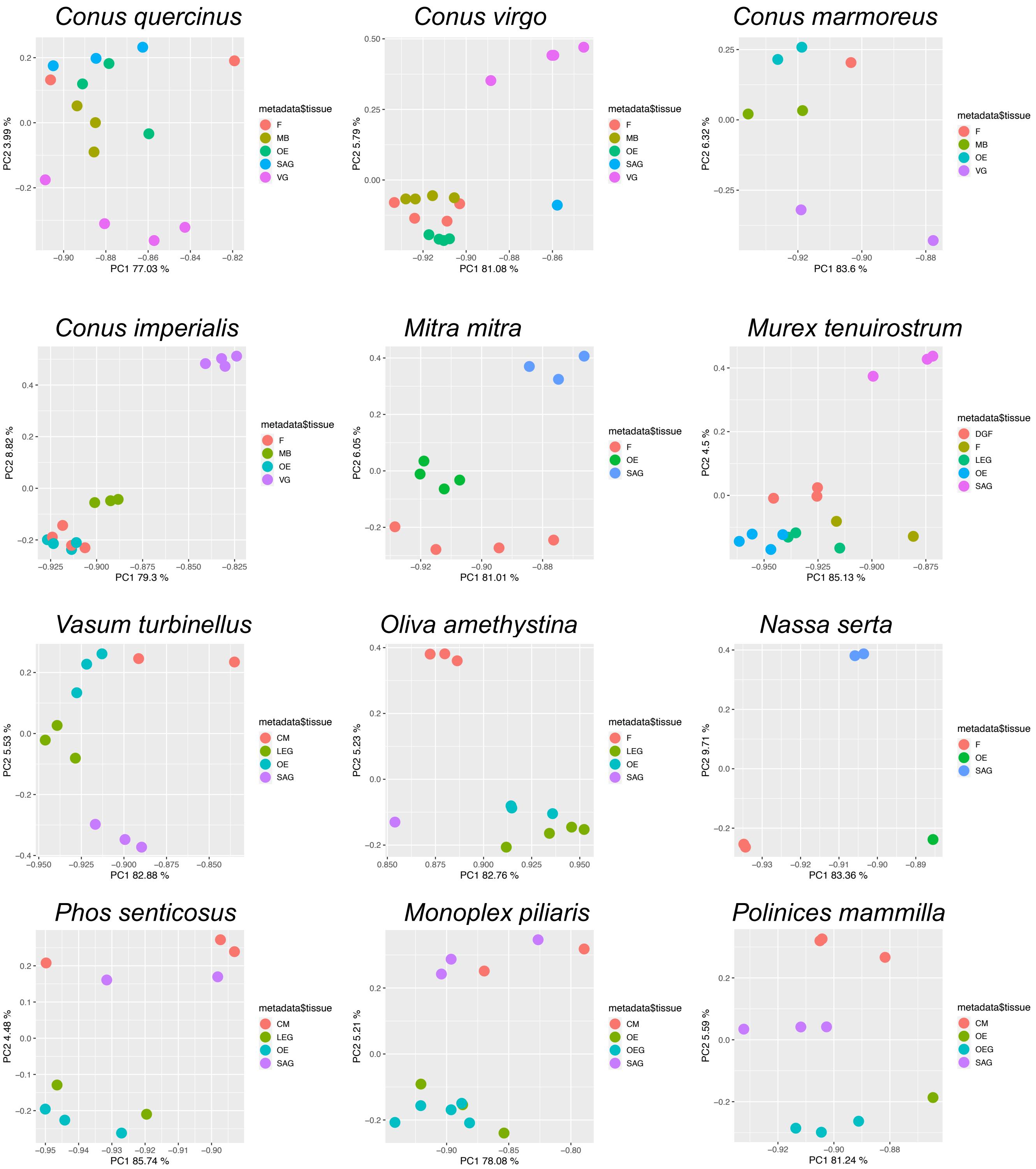


**Fig. S1. Principal component analysis (PCA) plot for each species individually after the quality-filtering step.** Each plot shows the variation along the first principal (PC1, x-axis) and the second principal component (PC2, y-axis). Samples are colored by tissue type.Tissue abbreviations: CM = columellar muscle; DGF = dorsal glandular folds; F = foot; LEG = gland of Leiblein; MB = muscular bulb; OE = oesophagus; OEG = oesophageal gland; SAG = salivary glands; VG = venom gland.

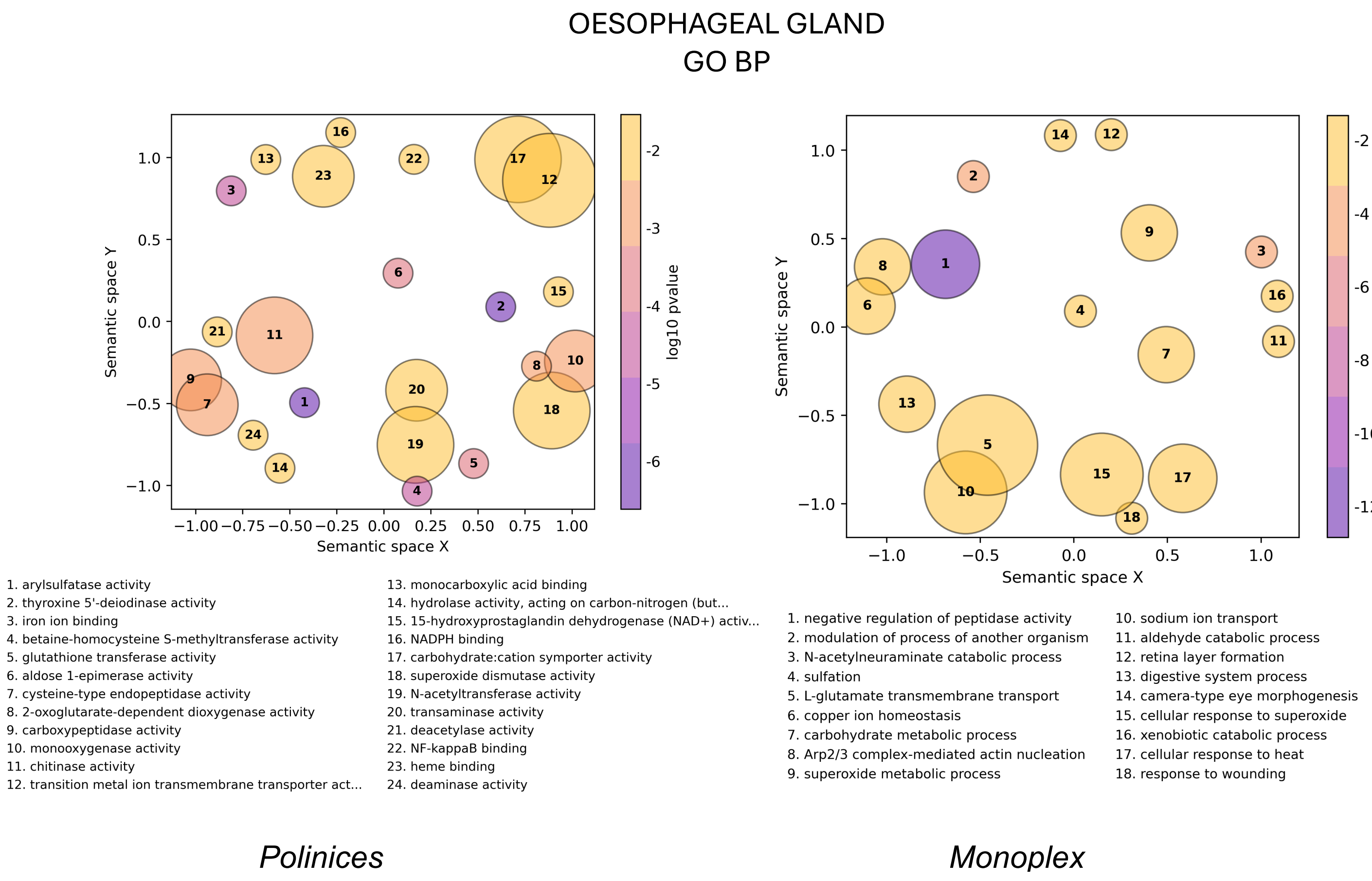


**Fig. S2. Gene Ontology biological process enrichment results for the oesophageal glands.** Each bubble represents a cluster of similar GO terms summarized by a representative term reported in the legend and sorted by the GO term p-values. Bubble size indicates the amount of GO terms in each cluster, and the color is the average p-value. Similar clusters plot closer to each other.

OESOPHAGEAL GLAND
GO MF

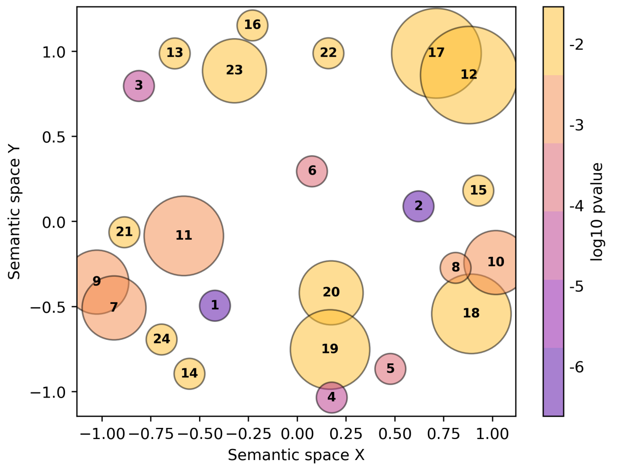


1. arylsulfatase activity
2. thyroxine 5'-deiodinase activity
3. iron ion binding
4. betaine-homocysteine S-methyltransferase activity
5. glutathione transferase activity
6. aldose 1-epimerase activity
7. cysteine-type endopeptidase activity
8. 2-oxoglutarate-dependent dioxygenase activity
9. carboxypeptidase activity
10. monooxygenase activity
11. chitinase activity
12. transition metal ion transmembrane transporter act...
13. monocarboxylic acid binding
14. hydrolase activity, acting on carbon-nitrogen (but...
15. 15-hydroxyprostaglandin dehydrogenase (NAD+) activ.
16. NADPH binding
17. carbohydrate:cation symporter activity
18. superoxide dismutase activity
19. N-acetyltransferase activity
20. transaminase activity
21. deacetylase activity
22. NF-kappaB binding
23. heme binding
24. deaminase activity

*Polinices mammilla*

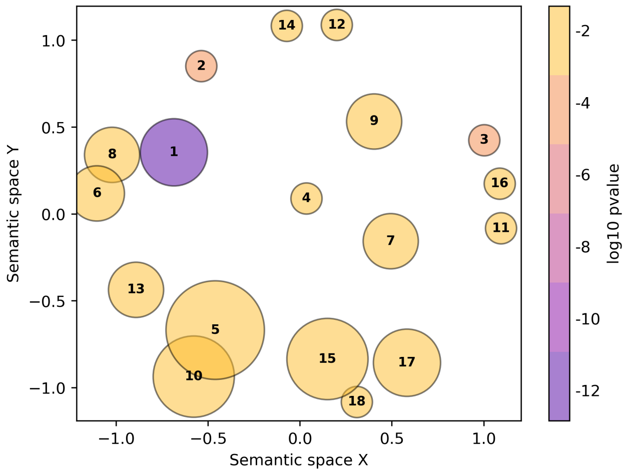


1. negative regulation of peptidase activity
2. modulation of process of another organism
3. N-acetylneuraminate catabolic process
4. sulfation
5. L-glutamate transmembrane transport
6. copper ion homeostasis
7. carbohydrate metabolic process
8. Arp2/3 complex-mediated actin nucleation
9. superoxide metabolic process
10. sodium ion transport
11. aldehyde catabolic process
12. retina layer formation
13. digestive system process
14. camera-type eye morphogenesis
15. cellular response to superoxide
16. xenobiotic catabolic process
17. cellular response to heat
18. response to wounding

*Monoplex piliaris*

**Fig. S3. Gene Ontology molecular function enrichment results for the oesophageal glands.** Each bubble represents a cluster of similar GO terms summarized by a representative term reported in the legend and sorted by the GO term p-values. Bubble size indicates the amount of GO terms in each cluster, and the color is the average p-value. Similar clusters plot closer to each other.

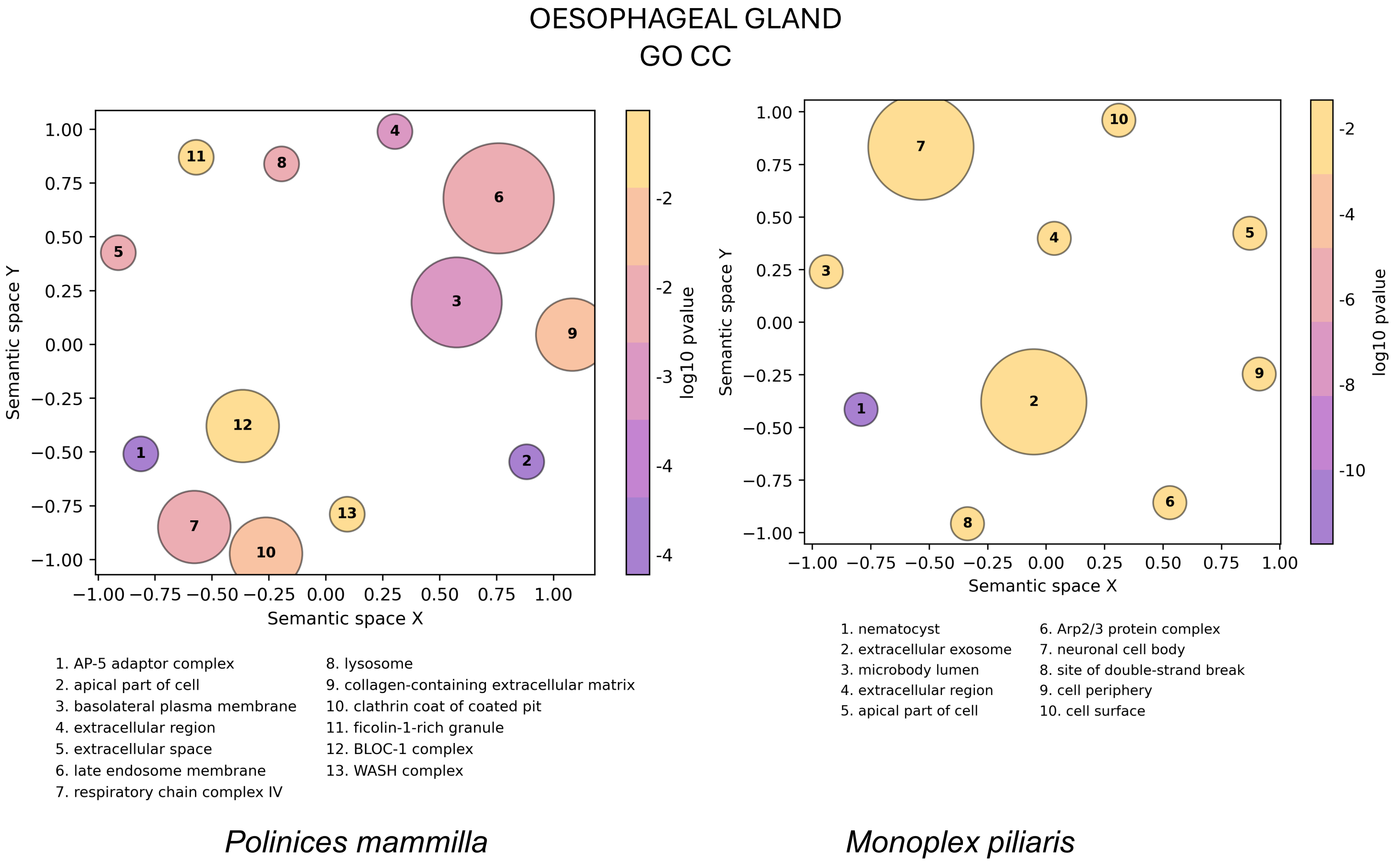


**Fig. S4. Gene Ontology cellular component enrichment results for the oesophageal glands.** Each bubble represents a cluster of similar GO terms summarized by a representative term reported in the legend and sorted by the GO term p-values. Bubble size indicates the amount of GO terms in each cluster, and the color is the average p-value. Similar clusters plot closer to each other.

GLAND OF LEIBLEIN
GO BP

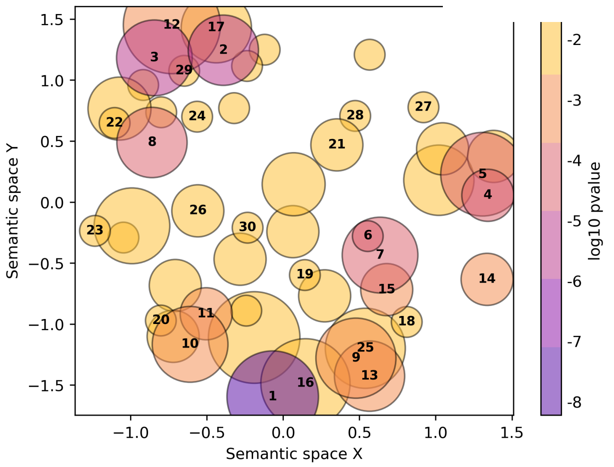


1. lysine catabolic process
2. succinate transmembrane transport
3. sodium ion transport
4. regulation of Arp2/3 complex-mediated actin nuclea...
5. cellular sodium ion homeostasis
6. sulfation
7. S-adenosylmethionine biosynthetic process
8. Golgi to plasma membrane transport
9. xenobiotic catabolic process
10. glutamate metabolic process
11. cobalamin metabolic process
12. carbohydrate transport
13. cellular carbohydrate catabolic process
14. neurotransmitter catabolic process
15. oligosaccharide biosynthetic process
16. polyol catabolic process
17. D-amino acid transport
18. collagen catabolic process
19. fructose 1,6-bisphosphate metabolic process
20. D-amino acid metabolic process
21. circadian sleep/wake cycle
22. import across plasma membrane
23. detection of chemical stimulus involved in ser
24. establishment or maintenance of transmembı
25. galactosylceramide catabolic process
26. phagosome-lysosome fusion involved in apop
27. regulation of dopamine metabolic process
28. copulation
29. import into cell
30. toxin metabolic process

*Murex tenuirostrum*

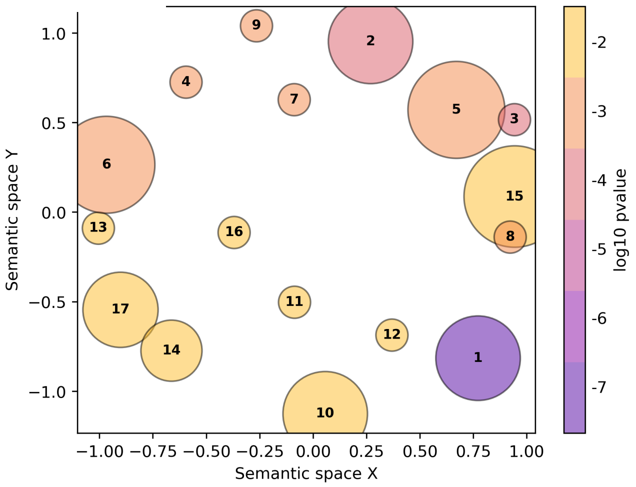


1. modulation of process of another organism
2. maintenance of location in cell
3. iron ion transport
4. thyroid hormone generation
5. L-glutamate transmembrane transport
6. peptidoglycan catabolic process
7. positive regulation of pigment cell differentiatio...
8. endosome to melanosome transport
9. negative regulation of molecular function
10. neuromuscular process controlling balance
11. bioluminescence
12. lysosome organization
13. proteolysis
14. water-soluble vitamin biosynthetic process
15. melanosome transport
16. aerobic respiration
17. methionine biosynthetic process

*Oliva amethystina*

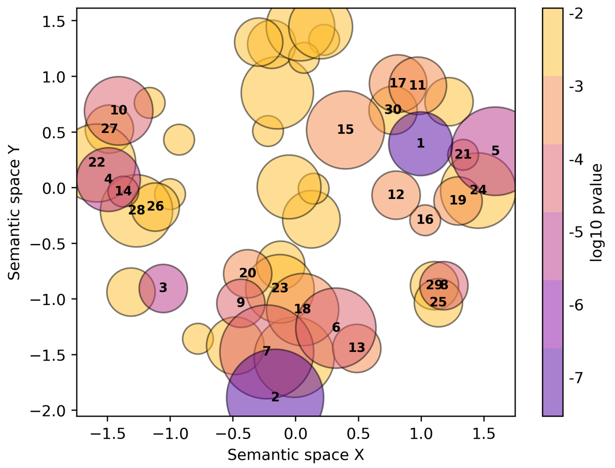


1. carbohydrate metabolic process
2. Arp2/3 complex-mediated actin nucleation
3. sequestering of metal ion
4. iron ion transport
5. monocarboxylic acid catabolic process
6. thyroid hormone generation
7. cellular iron ion homeostasis
8. decidualization
9. positive regulation of aspartic-type peptidase act...
10. quaternary ammonium group transport
11. dicarboxylic acid metabolic process
12. steroid metabolic process
13. neurotransmitter catabolic process
14. import into cell
15. removal of superoxide radicals
16. collagen catabolic process
17. L-ascorbic acid biosynthetic process
18. positive regulation of early endosome to late
19. heparan sulfate proteoglycan catabolic proce
20. regulation of cardiac muscle cell apoptotic pı
21. proteolysis
22. modified amino acid transport
23. regulation of superoxide metabolic process
24. ceramide catabolic process
25. bone development
26. melanosome transport
27. amide transport
28. endocytic recycling
29. endothelial cell development
30. glycoside catabolic process

*Vasum turbinellus*

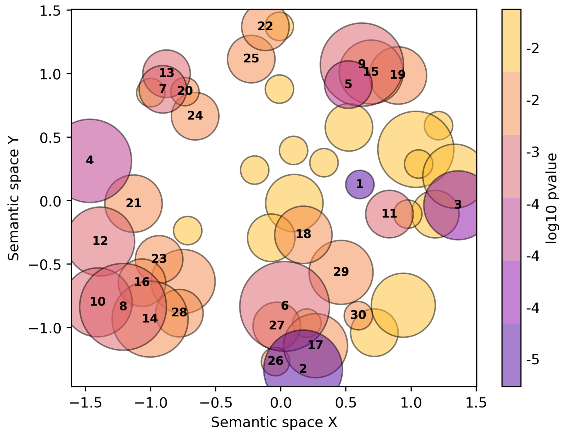


1. tricarboxylic acid cycle
2. sodium ion transport
3. gluconeogenesis
4. regulation of Arp2/3 complex-mediated actin nuclea...
5. glycoside catabolic process
6. endosomal transport
7. mitochondrion disassembly
8. regulation of metal ion transport
9. hydrogen peroxide catabolic process
10. positive regulation of cation channel activity
11. malate metabolic process
12. positive regulation of lamellipodium organization
13. vesicle organization
14. positive regulation of hemostasis
15. antibiotic catabolic process
16. negative regulation of NF-kappaB transcription fac...
17. L-glutamate transmembrane transport
18. ameboidal-type cell migration
19. ceramide catabolic process
20. vesicle tethering
21. negative regulation of T cell differentiation
22. thymus development
23. regulation of lipopolysaccharide-mediated signalin...
24. synaptic vesicle membrane organization
25. larval midgut cell programmed cell death
26. import into cell
27. export across plasma membrane
28. regulation of animal organ morphogenesis
29. cellular response to toxic substance
30. response to tumor cell

*Phos senticosus*

**Fig. S5. Gene Ontology biological process enrichment results for the glands of Leiblein.** Each bubble represents a cluster of similar GO terms summarized by a representative term reported in the legend and sorted by the GO term p-values. Bubble size indicates the amount of GO terms in each cluster, and the color is the average p-value. Similar clusters plot closer to each other.

### GLAND OF LEIBLEIN
## GO MF

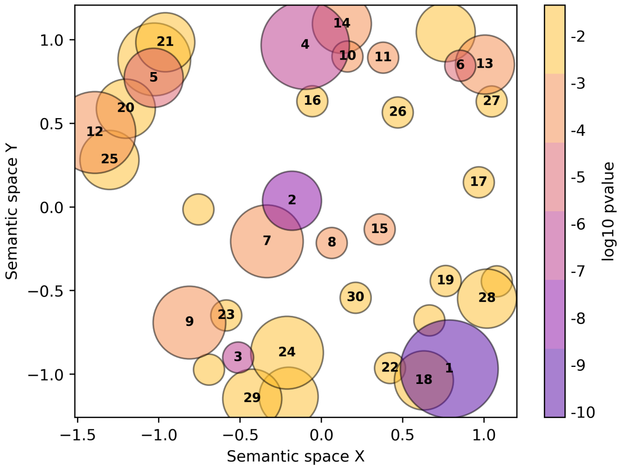


1. aminopeptidase activity
2. hydroxylysine kinase activity
3. ferroxidase activity
4. ferric iron binding
5. succinate transmembrane transporter activity
6. proteoglycan binding
7. aryl sulfotransferase activity
8. S-methyltransferase activity
9. D-amino-acid oxidase activity
10. FAD binding
11. heme binding
12. solute:sodium symporter activity
13. clathrin binding
14. phosphatidylinositol binding
15. methionine adenosyltransferase activity
16. collagen binding
17. cysteine-type endopeptidase inhibitor activity
18. omega peptidase activity
19. gluconolactonase activity
20. transmembrane transporter activity
21. glucose transmembrane transporter activity
22. hydrolase activity, acting on carbon-nitrogen (but...
23. 15-oxoprostaglandin 13-oxidase activity
24. 13-prostaglandin reductase activity
25. P-type sodium transporter activity
26. quinone binding
27. fibronectin binding
28. galactosylceramidase activity
29. oxidoreductase activity, acting on the aldehyde or...
30. pyruvate dehydrogenase (acetyl-transferring) kinas...

*Murex tenuirostrum*

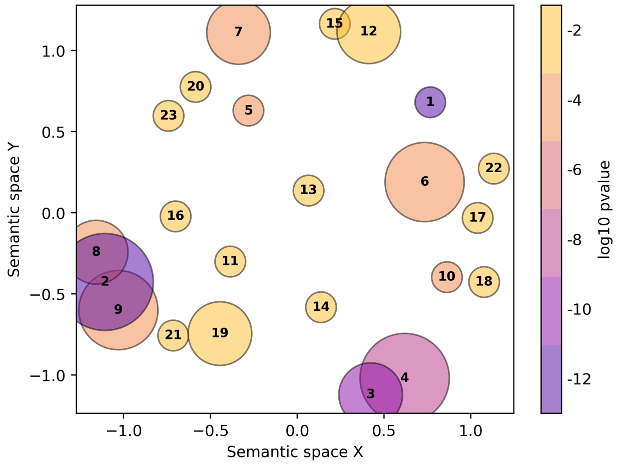


1. toxin activity
2. aminopeptidase activity
3. acetylcholine receptor inhibitor activity
4. ion channel inhibitor activity
5. ferroxidase activity
6. ferric iron binding
7. cytochrome-c oxidase activity
8. cysteine-type peptidase activity
9. metalloendopeptidase activity
10. chitin binding
11. gluconolactonase activity
12. glucose:sodium symporter activity
13. peptidoglycan muralytic activity
14. betaine-homocysteine S-methyltransferase activity
15. succinate transmembrane transporter activity
16. bis(5'-adenosyl)-triphosphatase activity
17. collagen binding
18. proteoglycan binding
19. beta-N-acetylhexosaminidase activity
20. 2-oxoglutarate-dependent dioxygenase activity
21. hydrolase activity, acting on carbon-nitrogen (but...
22. peptidoglycan binding
23. 15-oxoprostaglandin 13-oxidase activity

*Oliva amethystina*

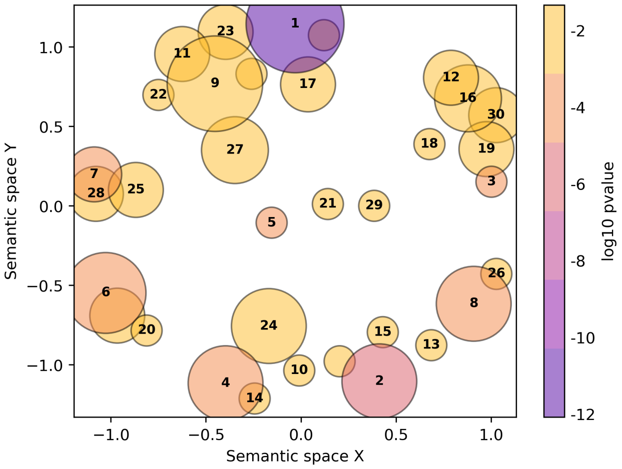


1. aminopeptidase activity
2. ferrous iron binding
3. ferroxidase activity
4. actin binding
5. betaine-homocysteine S-methyltransferase activity
6. L-amino acid transmembrane transporter activity
7. L-fuconate dehydratase activity
8. ornithine decarboxylase regulator activity
9. alpha-amylase activity
10. heme binding
11. acid phosphatase activity
12. oxidoreductase activity, acting on paired donors, ...
13. collagen binding
14. proteoglycan binding
15. phosphatidylinositol binding
16. oxidoreductase activity, acting on the CH-NH2
17. hydrolase activity, acting on ether bonds
18. 2-alkenal reductase [NAD(P)+] activity
19. superoxide dismutase activity
20. neurotransmitter transmembrane transporter
21. glutathione transferase activity
22. gluconolactonase activity
23. sulfuric ester hydrolase activity
24. folic acid binding
25. oxo-acid-lyase activity
26. structural constituent of eye lens
27. beta-ureidopropionase activity
28. aconitate hydratase activity
29. transketolase or transaldolase activity
30. monooxygenase activity

*Vasum turbinellus*

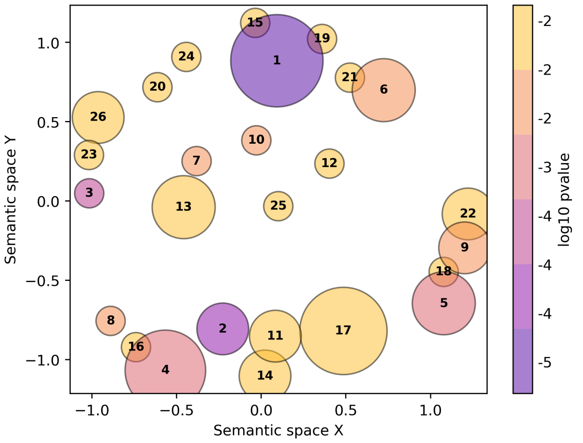


1. peptidyl-dipeptidase activity
2. phosphatidylinositol binding
3. L-malate dehydrogenase activity
4. clathrin adaptor activity
5. proton-transporting ATPase activity, rotational me...
6. alpha-L-fucosidase activity
7. 1-phosphatidylinositol-4-phosphate 5-kinase activi...
8. proteoglycan binding
9. symporter activity
10. S-methyltransferase activity
11. transition metal ion binding
12. C-acyltransferase activity
13. glycoprotein-N-acetylgalactosamine 3-beta-galactos...
14. heme binding
15. hydrolase activity, acting on carbon-nitrogen (but...
16. protease binding
17. GDP-dissociation inhibitor activity
18. wide pore channel activity
19. hydrolase activity, acting on ether bonds
20. hydro-lyase activity
21. arylsulfatase activity
22. nucleoside transmembrane transporter activity
23. peroxiredoxin activity
24. catalytic activity
25. aryl sulfotransferase activity
26. oxidoreductase activity, acting on paired donors, ...

*Phos senticosus*

**Fig. S6. Gene Ontology molecular function enrichment results for the glands of Leiblein.** Each bubble represents a cluster of similar GO terms summarized by a representative term reported in the legend and sorted by the GO term p-values. Bubble size indicates the amount of GO terms in each cluster, and the color is the average p-value. Similar clusters plot closer to each other.

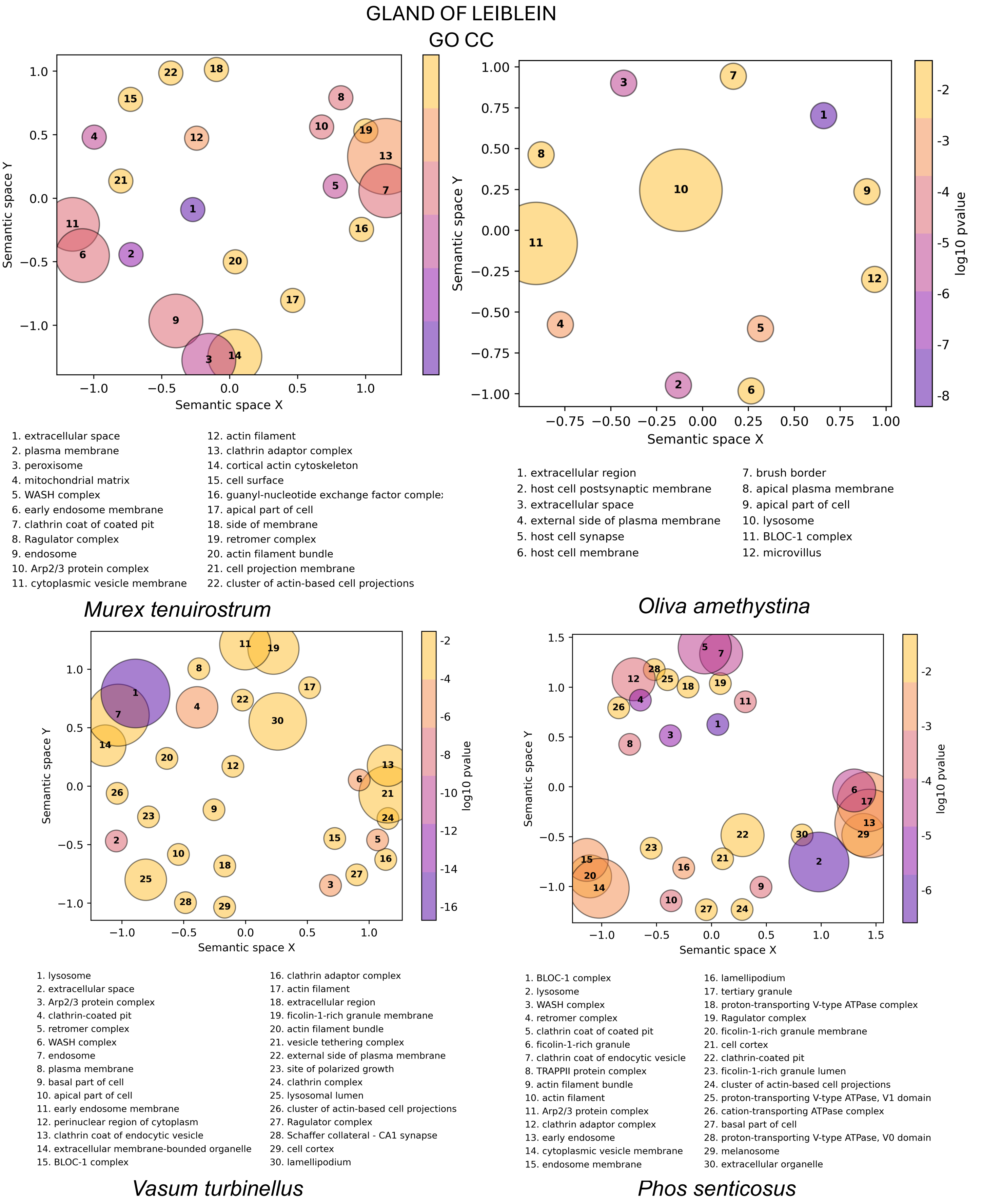


**Fig. S7. Gene Ontology cellular component enrichment results for the glands of Leiblein.** Each bubble represents a cluster of similar GO terms summarized by a representative term reported in the legend and sorted by the GO term p-values. Bubble size indicates the amount of GO terms in each cluster, and the color is the average p-value. Similar clusters plot closer to each other.

### VENOM GLAND
## GO BP

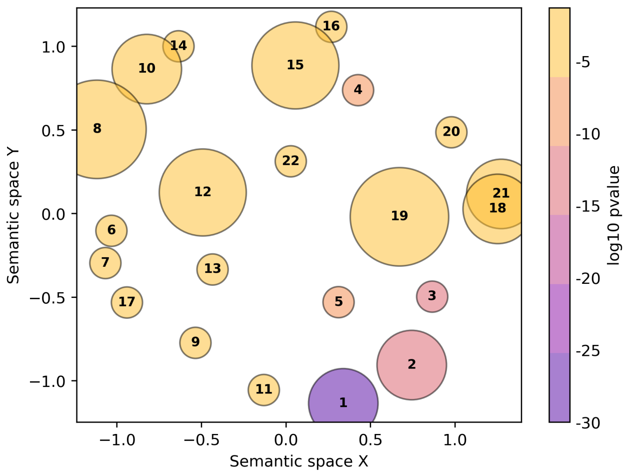


1. modulation of process of another organism
2. positive regulation of voltage-gated sodium channe...
3. negative regulation of signaling receptor activity
4. synaptic transmission, cholinergic
5. chemical synaptic transmission, postsynaptic
6. peptidyl-glutamic acid carboxylation
7. peptidyl-proline hydroxylation
8. ubiquitin-dependent ERAD pathway
9. reverse transcription involved in RNA-mediated tra...
10. response to vitamin K
11. lung growth
12. ornithine metabolic process
13. RNA phosphodiester bond hydrolysis, exonucleolyti
14. response to manganese ion
15. establishment of protein localization to endoplasm
16. transition metal ion transport
17. signal peptide processing
18. cellular zinc ion homeostasis
19. negative regulation of microtubule depolymerizatic
20. sequestering of metal ion
21. cellular transition metal ion homeostasis
22. microtubule depolymerization

*Conus quercinus*

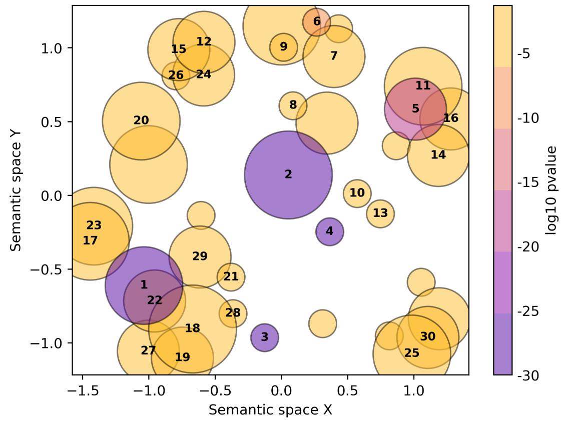


1. negative regulation of signaling receptor activity
2. modulation of receptor activity in another organis...
3. chemical synaptic transmission, postsynaptic
4. synaptic transmission, cholinergic
5. sodium ion transmembrane transport
6. peptidyl-proline hydroxylation
7. signal peptide processing
8. spermine biosynthetic process
9. ornithine metabolic process
10. Golgi organization
11. L-arginine transmembrane transport
12. response to endoplasmic reticulum stress
13. endoplasmic reticulum membrane organization
14. establishment of protein localization to endoplasm...
15. response to unfolded protein
16. post-translational protein targeting to membrane, ...
17. positive regulation of nitric oxide biosynthetic p...
18. negative regulation of T cell differentiation
19. negative regulation of lipopolysaccharide-mediated...
20. regulation of translational initiation by eIF2 alp...
21. regulation of systemic arterial blood pressure
22. regulation of extrinsic apoptotic signaling pathwa...
23. regulation of nitric oxide biosynthetic process
24. natural killer cell mediated cytotoxicity
25. labyrinthine layer development
26. response to tumor cell
27. positive regulation of blood coagulation
28. extrinsic apoptotic signaling pathway in absence o...
29. lipopolysaccharide-mediated signaling pathway
30. embryonic placenta development

*Conus virgo*

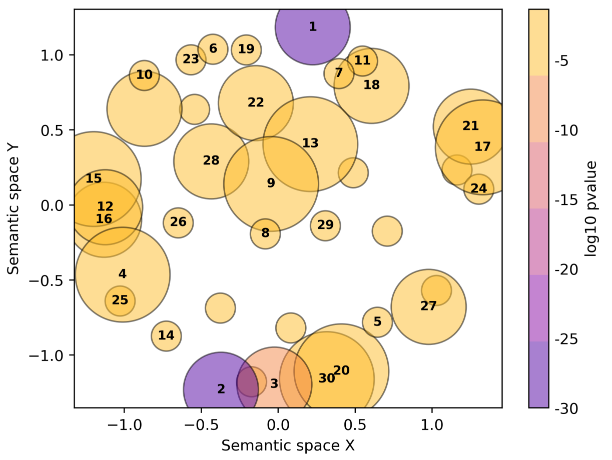


1. modulation of process of another organism
2. positive regulation of voltage-gated sodium channe...
3. negative regulation of signaling receptor activity
4. response to endoplasmic reticulum stress
5. chemical synaptic transmission, postsynaptic
6. signal peptide processing
7. endoplasmic reticulum membrane organization
8. synaptic transmission, cholinergic
9. protein folding
10. RNA-templated DNA biosynthetic process
11. Golgi organization
12. endoplasmic reticulum to Golgi vesicle-mediated tr...
13. transposition
14. SREBP signaling pathway
15. response to unfolded protein
16. establishment of protein localization to endoplasm...
17. placenta development
18. extracellular matrix organization
19. protein deglutathionylation
20. negative regulation of protein localization to pla...
21. maternal placenta development
22. ornithine metabolic process
23. peptidyl-proline hydroxylation
24. somite development
25. inflammatory response
26. amino acid transmembrane transport
27. embryonic axis specification
28. S-adenosylmethionine biosynthetic process
29. macrophage activation
30. regulation of endothelial cell migration

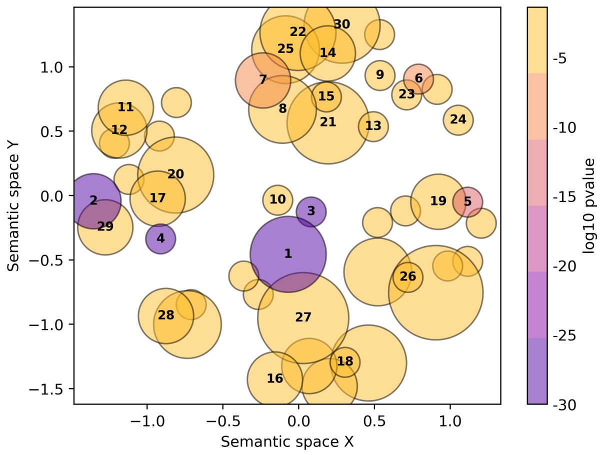


1. modulation of receptor activity in another organis...
2. negative regulation of signaling receptor activity
3. synaptic transmission, cholinergic
4. chemical synaptic transmission, postsynaptic
5. sodium ion transmembrane transport
6. peptidyl-proline hydroxylation
7. one-carbon metabolic process
8. S-adenosylmethionine biosynthetic process
9. signal peptide processing
10. hemolysis in another organism
11. G protein-coupled receptor signaling pathway
12. endoplasmic reticulum unfolded protein response
13. reverse transcription involved in RNA-mediated tra...
14. purine ribonucleoside catabolic process
15. RNA phosphodiester bond hydrolysis, exonucleolytic
16. nematode larval development
17. regulation of post-embryonic development
18. cuticle development involved in collagen and cutic...
19. retrograde vesicle-mediated transport, Golgi to en...
20. positive regulation of receptor internalization
21. dopamine metabolic process
22. tetrahydrobiopterin biosynthetic process
23. protein ufmylation
24. cellular modified amino acid catabolic process
25. serine family amino acid biosynthetic process
26. pore complex assembly
27. oviposition
28. response to xenobiotic stimulus
29. negative regulation of response to endoplasmic ret...
30. IMP biosynthetic process

*Conus marmoreus*

*Conus imperialis*

**Fig. S8. Gene Ontology biological process enrichment results for the venom glands.** Each bubble represents a cluster of similar GO terms summarized by a representative term reported in the legend,and sorted by the GO term p-values. Bubble size indicates the amount of GO terms in each cluster, and the color is the average p-value. Similar clusters plot closer to each other.

VENOM GLAND
GO MF

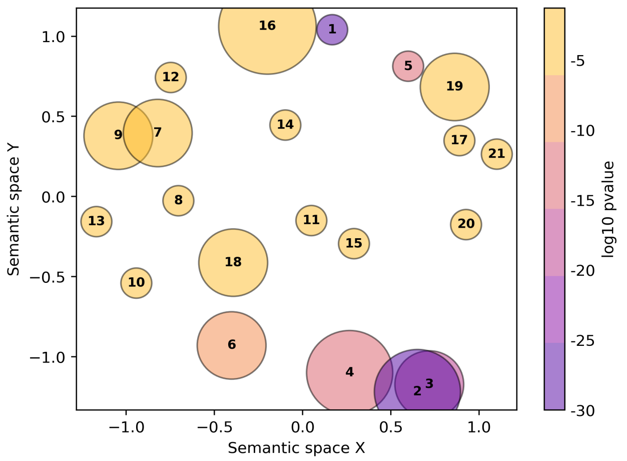


1. toxin activity
2. ion channel inhibitor activity
3. sodium channel regulator activity
4. acetylcholine receptor inhibitor activity
5. acetylcholine binding
6. acetylcholine receptor activity
7. peptidyl-dipeptidase activity
8. procollagen-proline 4-dioxygenase activity
9. metalloendopeptidase activity
10. RNA-directed DNA polymerase activity
11. amidine-lyase activity
12. exoribonuclease H activity
13. type II site-specific deoxyribonuclease activity
14. intramolecular oxidoreductase activity, transposin...
15. gamma-glutamyl carboxylase activity
16. sodium channel activity
17. L-ascorbic acid binding
18. ferroxidase activity
19. ferric iron binding
20. transmembrane transporter binding
21. flavin adenine dinucleotide binding

*Conus quercinus*

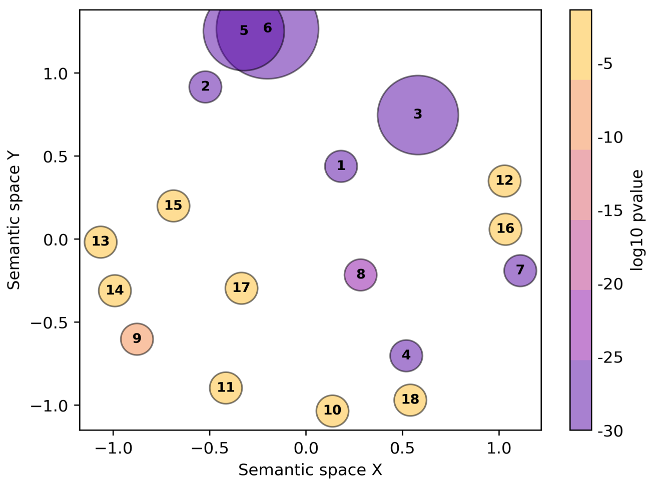


1. toxin activity
2. acetylcholine receptor inhibitor activity
3. acetylcholine receptor activity
4. acetylcholine binding
5. potassium channel regulator activity
6. sodium channel inhibitor activity
7. sodium channel activity
8. transmembrane transporter binding
9. procollagen-proline 4-dioxygenase activity
10. L-ascorbic acid binding
11. intramolecular oxidoreductase activity, transposin...
12. L-arginine transmembrane transporter activity
13. disulfide oxidoreductase activity
14. peroxiredoxin activity
15. calcium-dependent cysteine-type endopeptidase acti...
16. amide transmembrane transporter activity
17. amidine-lyase activity
18. iron ion binding

*Conus virgo*

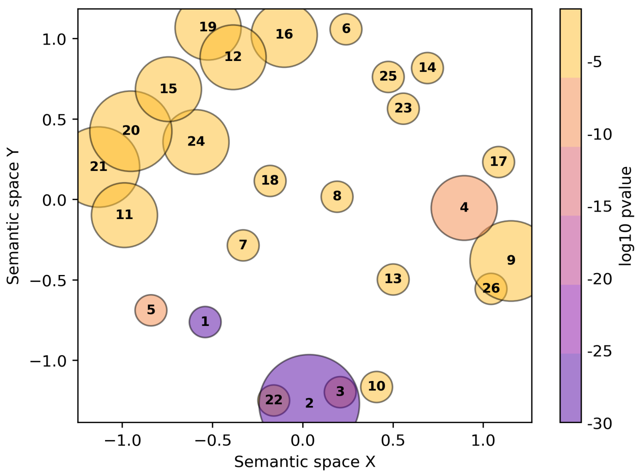


1. toxin activity
2. ion channel inhibitor activity
3. acetylcholine receptor inhibitor activity
4. acetylcholine receptor activity
5. acetylcholine binding
6. procollagen-proline 4-dioxygenase activity
7. intramolecular oxidoreductase activity, transposin...
8. amidine-lyase activity
9. sodium channel activity
10. peptidase activator activity involved in apoptotic...
11. RNA-directed DNA polymerase activity
12. peptidyl-dipeptidase activity
13. L-ascorbic acid binding
14. disulfide oxidoreductase activity
15. serine hydrolase activity
16. aminoacyltransferase activity
17. misfolded protein binding
18. methionine adenosyltransferase activity
19. aspartic-type endopeptidase activity
20. exoribonuclease H activity
21. type II site-specific deoxyribonuclease activity
22. cysteine-type endopeptidase regulator activity
23. malate dehydrogenase activity
24. alpha-L-fucosidase activity
25. peroxiredoxin activity
26. amino acid transmembrane transporter activity

*Conus marmoreus*

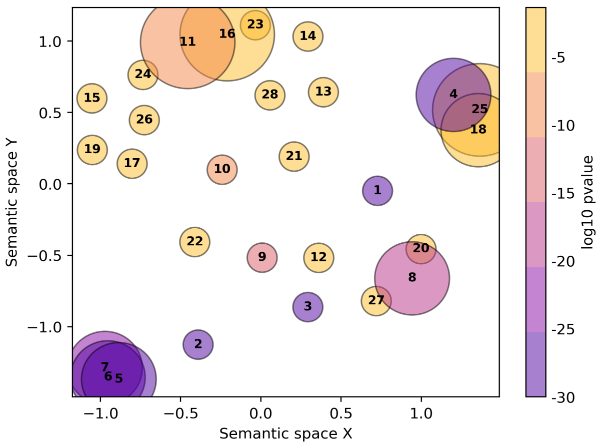


1. toxin activity
2. acetylcholine receptor inhibitor activity
3. acetylcholine binding
4. acetylcholine receptor activity
5. ion channel regulator activity
6. ion channel inhibitor activity
7. sodium channel inhibitor activity
8. sodium channel activity
9. transmembrane transporter binding
10. intramolecular oxidoreductase activity, transposin...
11. procollagen-proline 4-dioxygenase activity
12. L-ascorbic acid binding
13. methionine adenosyltransferase activity
14. RNA-directed DNA polymerase activity
15. type II site-specific deoxyribonuclease activity
16. disulfide oxidoreductase activity
17. adenosylhomocysteinase activity
18. peptide receptor activity
19. exoribonuclease H activity
20. L-glutamate transmembrane transporter activity
21. GDP-mannose 4,6-dehydratase activity
22. peptide binding
23. oxidoreductase activity, acting on the CH-NH group...
24. glutathione hydrolase activity
25. G protein-coupled receptor activity
26. cyclohydrolase activity
27. membrane potential driven uniporter activity
28. hydroxymethyl-, formyl- and related transferase ac...

*Conus imperialis*

**Fig. S9. Gene Ontology molecular function enrichment results for the venom glands.** Each bubble represents a cluster of similar GO terms summarized by a representative term reported in the legend and sorted by the GO term p-values. Bubble size indicates the amount of GO terms in each cluster, and the color is the average p-value. Similar clusters plot closer to each other.

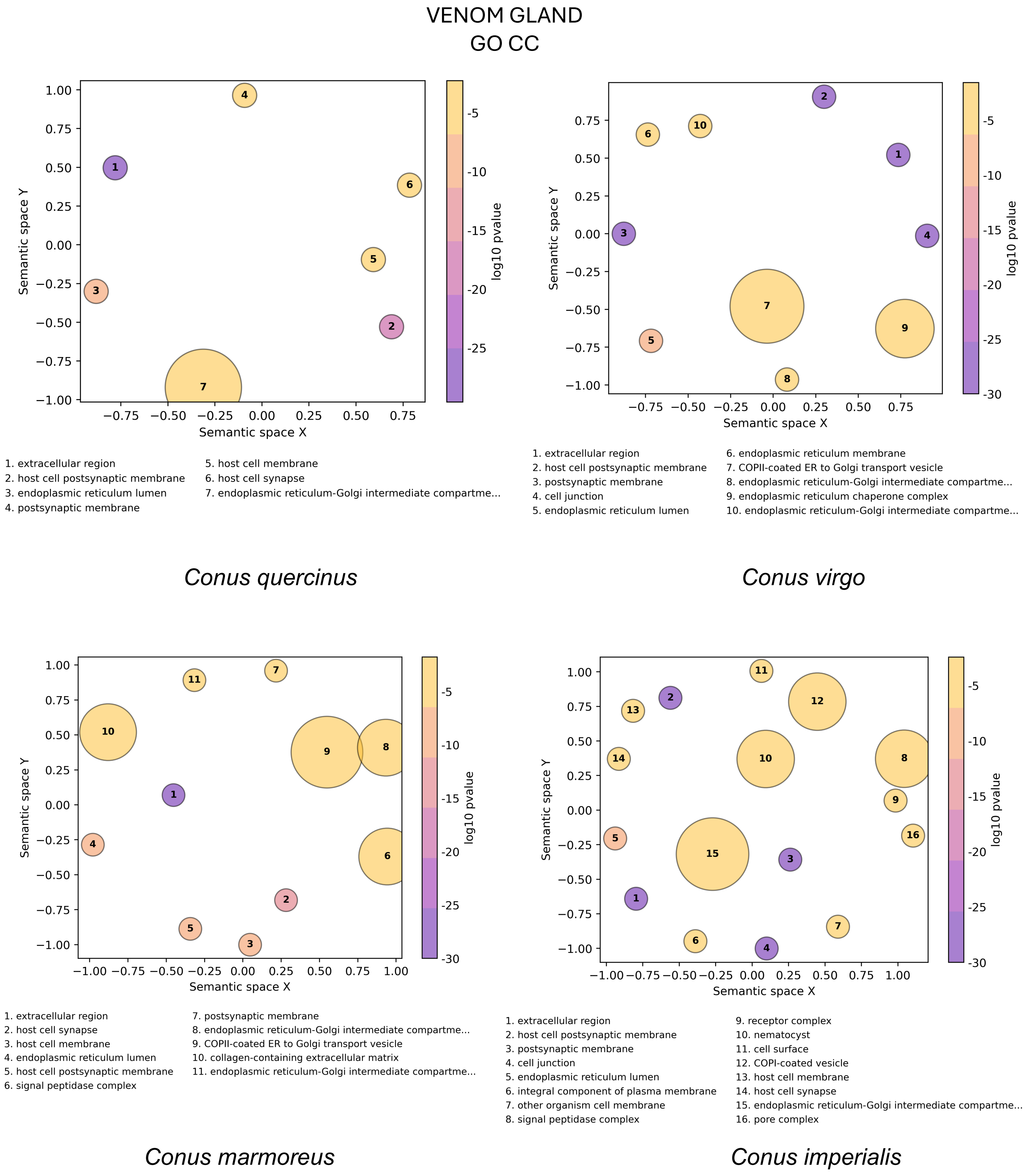


**Fig. S10. Gene Ontology cellular compartment enrichment results for the venom glands.** Each bubble represents a cluster of similar GO terms summarized by a representative term reported in the legend and sorted by the GO term p-values. Bubble size indicates the amount of GO terms in each cluster, and the color is the average p-value. Similar clusters plot closer to each other.

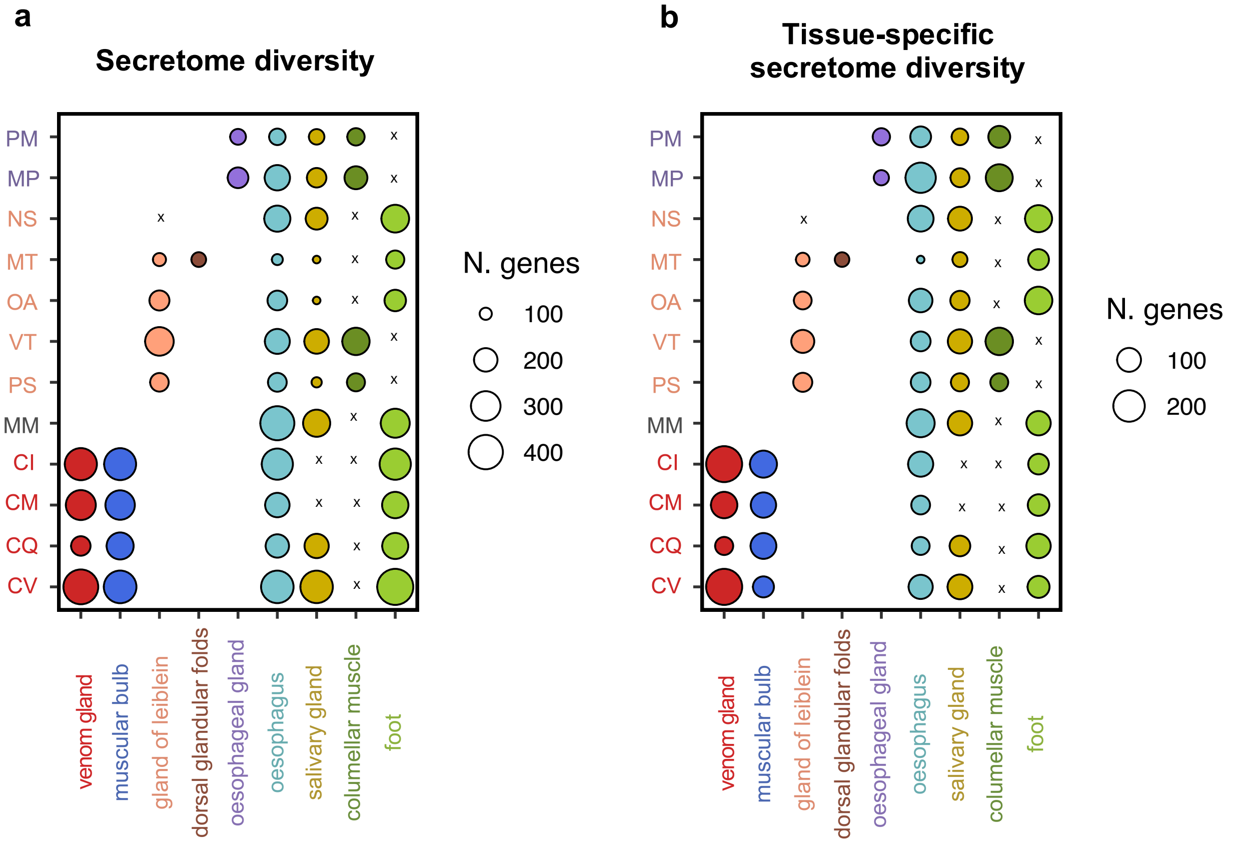


**Fig. S11. Secretome diversity.** a) Number of genes predicted to have a signal peptide in the whole transcriptome. The venom gland has a higher number (mean N = 565) compared to the gland of Leiblein (N = 354; t = 2, df = 5.8, p = 0.04) and the oesophageal gland (mean N = 379; t = 1, df = 1,3, p = 0.23), although the latter is not statistically significant. b) Number of tissue-specific genes with a signal peptide. The venom gland has a higher number of tissue-specific genes (mean N = 183) compared to the oesophageal gland (mean N = 38; t = 2.5, df = 3, p = 0.04) and the gland of Leiblein (mean N = 55; t = 2.2, df = 3.3, p = 0.05).

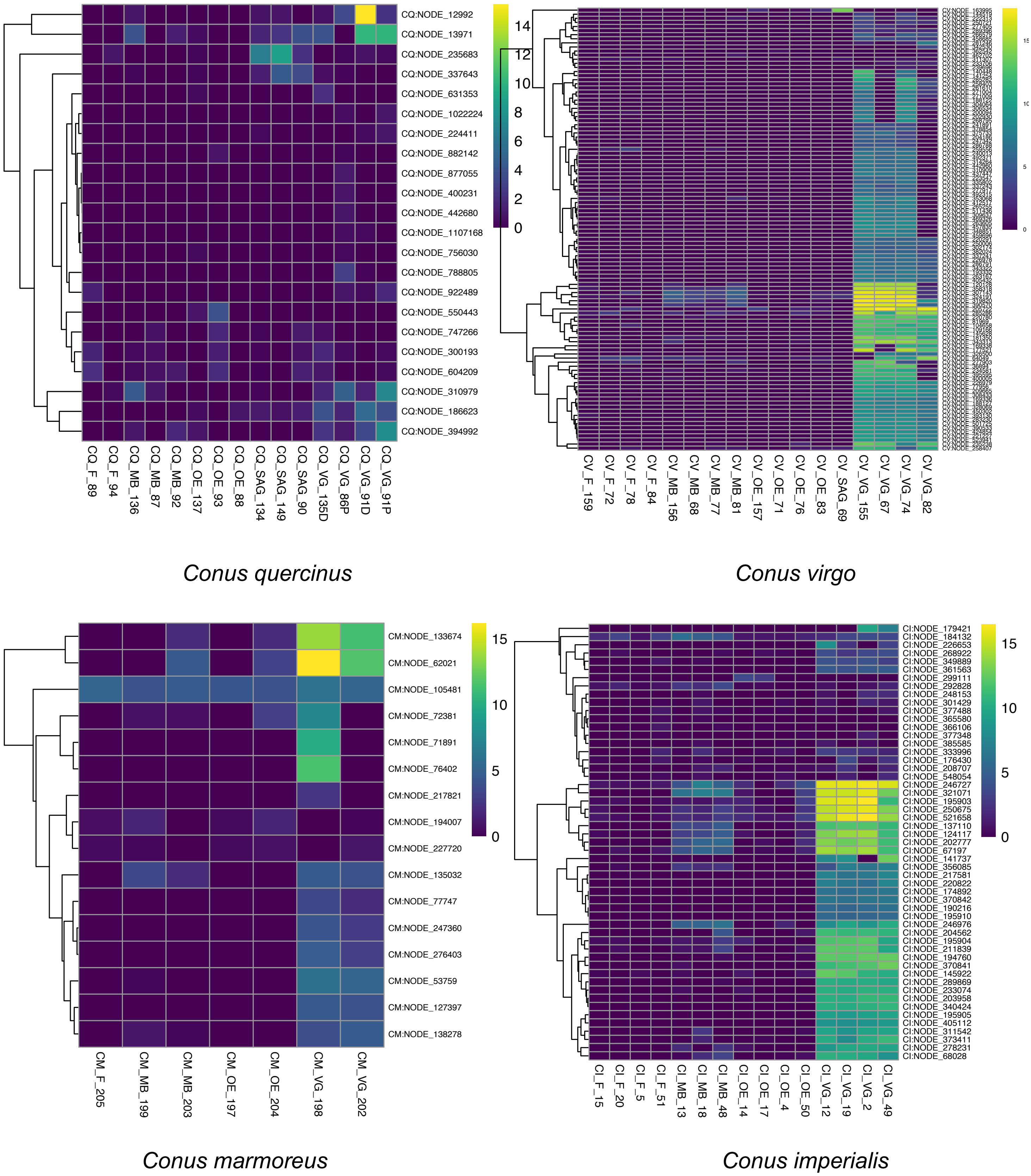


**Fig. S12. Expression profiles of conotoxins in venomous species.** Only transcripts with a predicted signal peptide and Pfam domain predicted as conotoxin are reported. Expression levels are in log2(TPM). VG = venom gland, MB = muscular venom bulb, SAG = salivary glands, OE = oesophagus, F = foot.

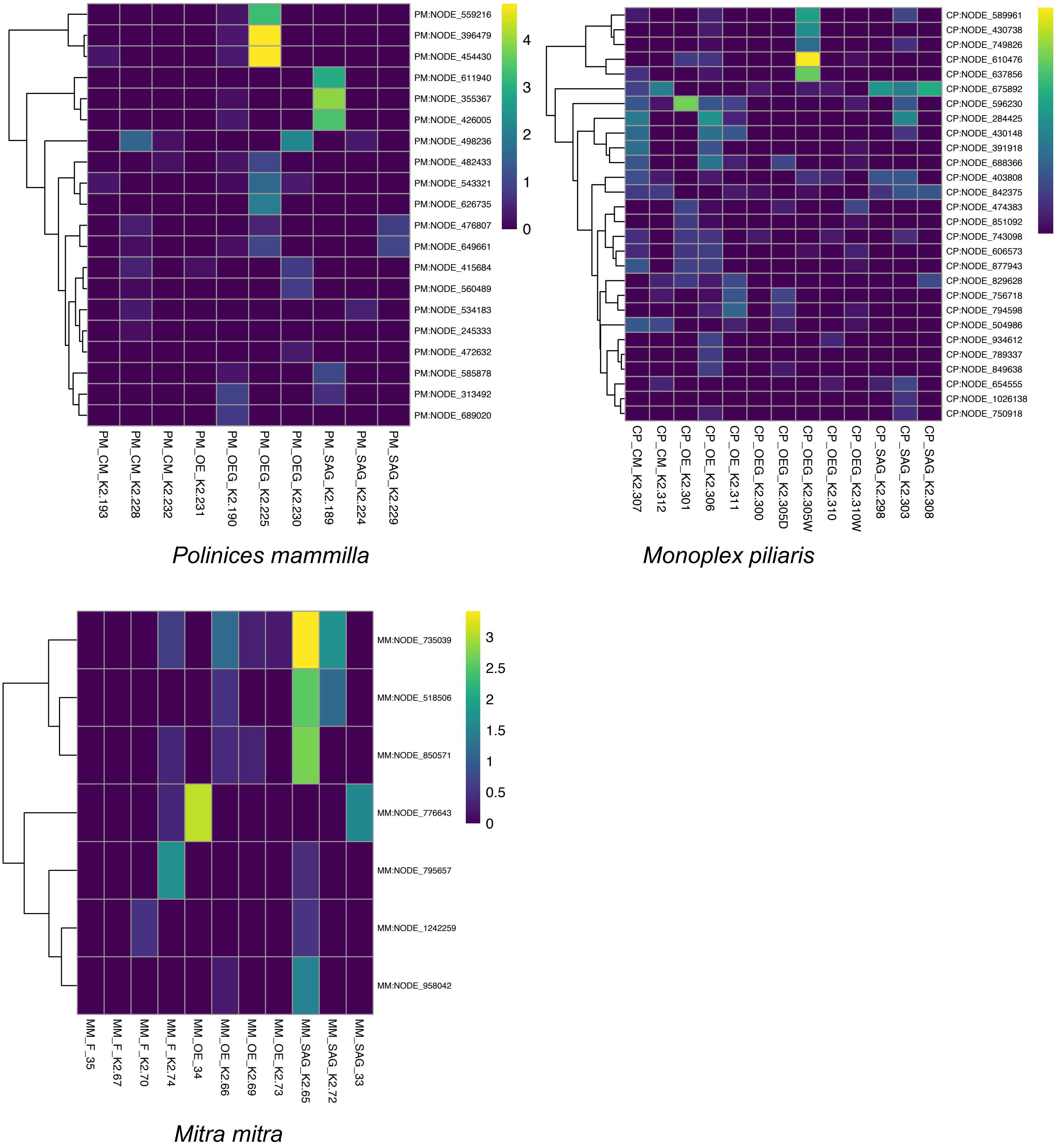


**Fig. S13. Expression profiles of conotoxins in species with oesophageal glands and the glandless species (*Mitra mitra*).** Only transcripts with a predicted signal peptide and Pfam domain predicted as conotoxin are reported. Expression levels are in log2(TPM). OEG = oesophageal gland, SAG = salivary glands, OE = oesophagus, F = foot, CM = columellar muscle.

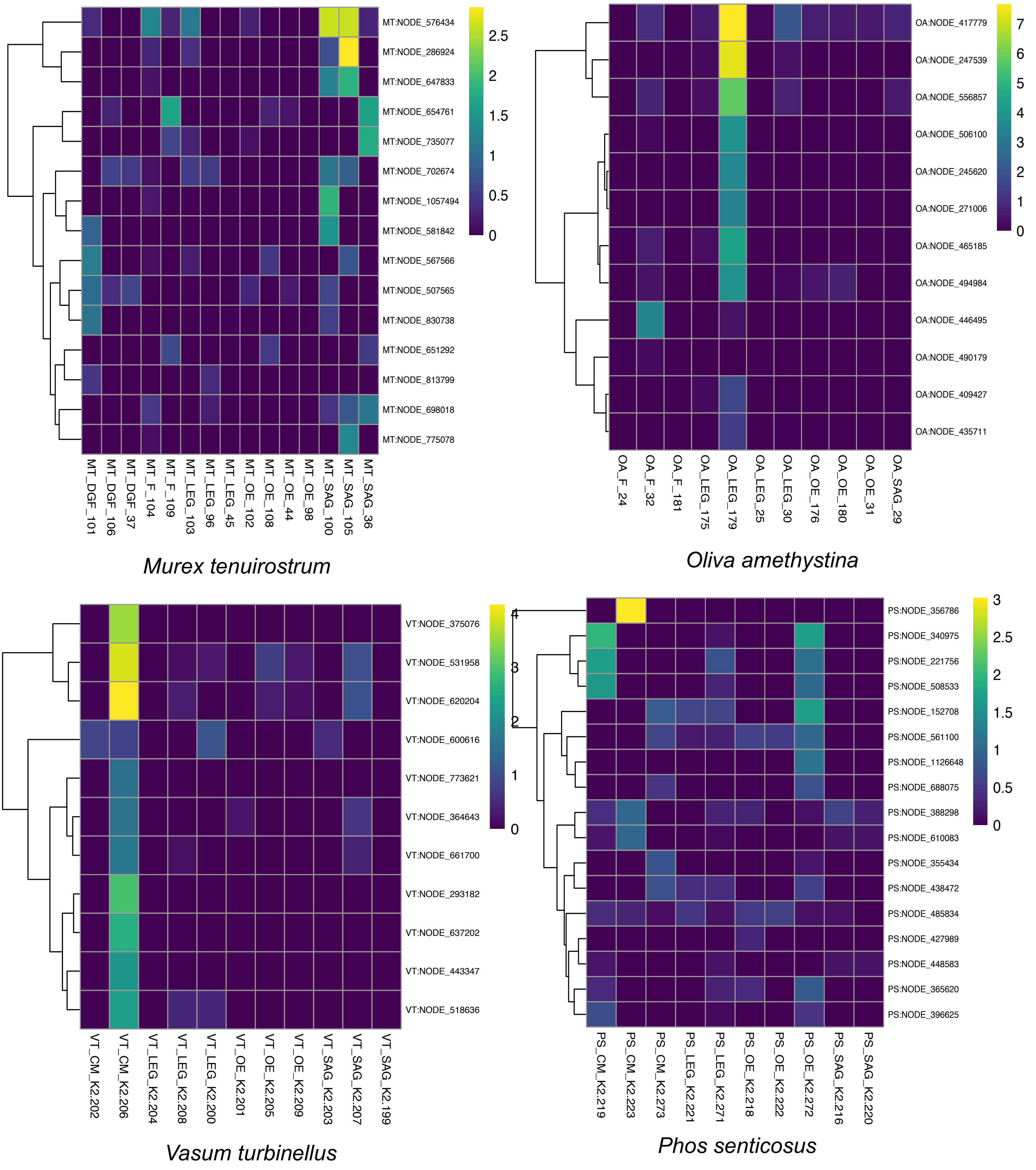


**Fig. S14. Expression profiles of conotoxins in species with the gland of Leiblein.** Only transcripts with a predicted signal peptide and Pfam domain predicted as conotoxin are reported. Expression levels are in log2(TPM). LEG = gland of Leiblein, SAG = salivary glands, OE = oesophagus, F = foot, CM = columellar muscle, DGF = dorsal glandular folds.

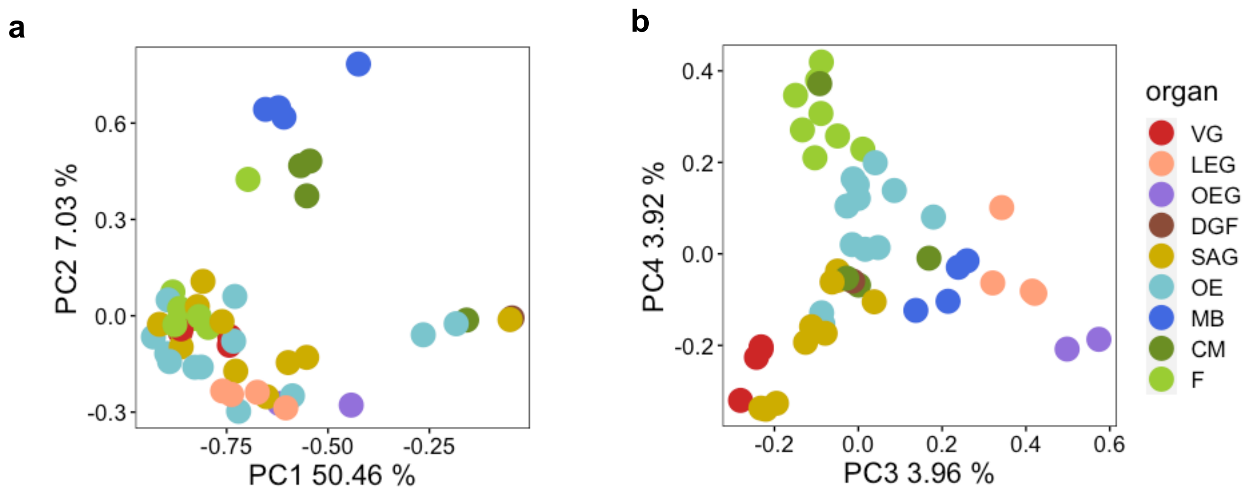


**Fig. S15. Principal component analysis plot using the random-based gene expression matrix.** a) First and second components. b) Third and fourth components. Tissue abbreviations as in previous figures.

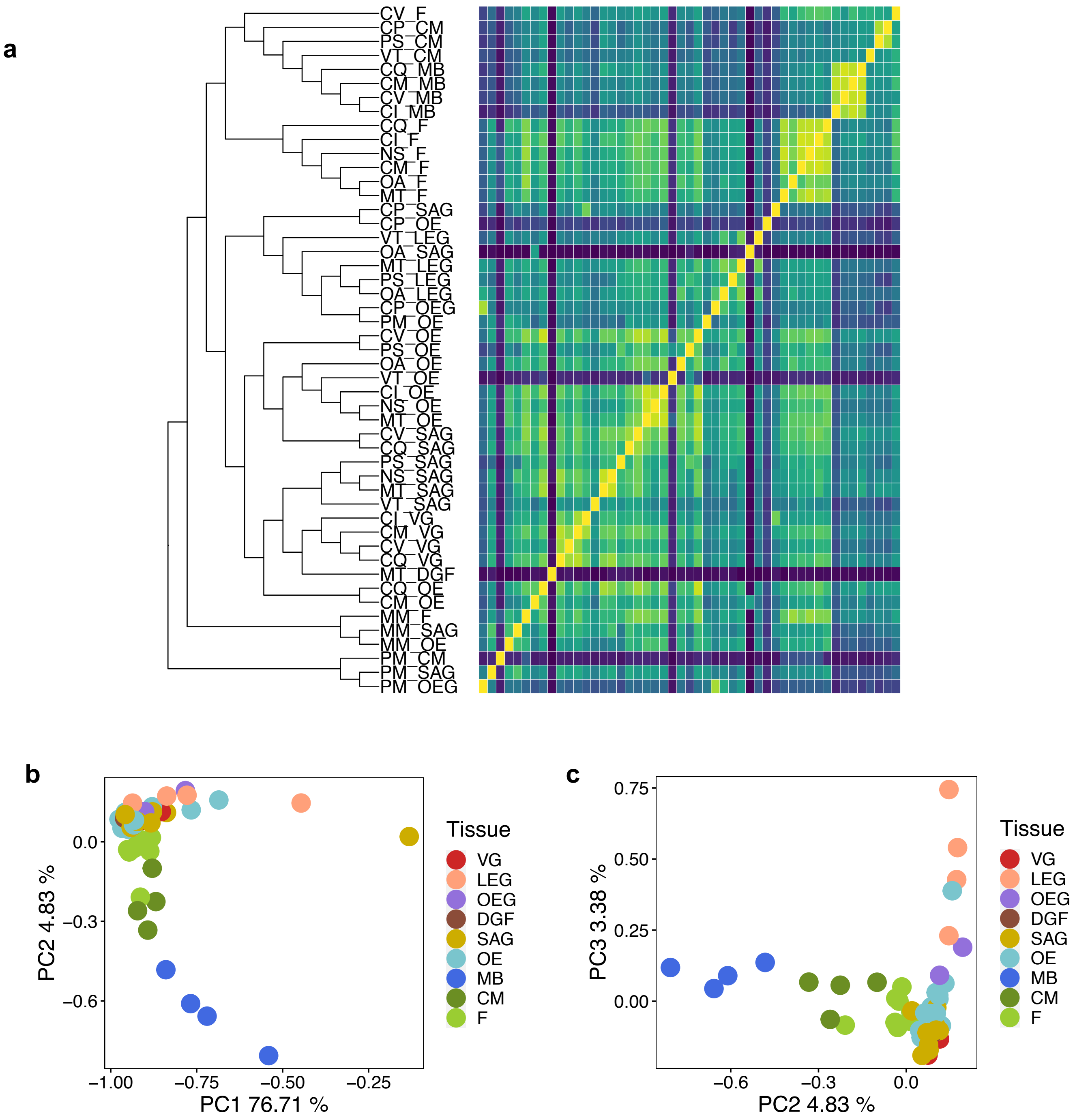


**Fig. S16. Transcriptome similarity and principal component analysis plot using the mean-based gene expression matrix.** a) Heatmap of Pearson correlation coefficients between tissues and species. The expression tree was made using neighbour-joining based on the correlation matrix. b) Principal component analysis plot with the first and second components, and c) third and fourth components. The orthogroup expression matrix used was based on the mean TPM across all the genes within an orthogroup. Abbreviations as in previous figures.

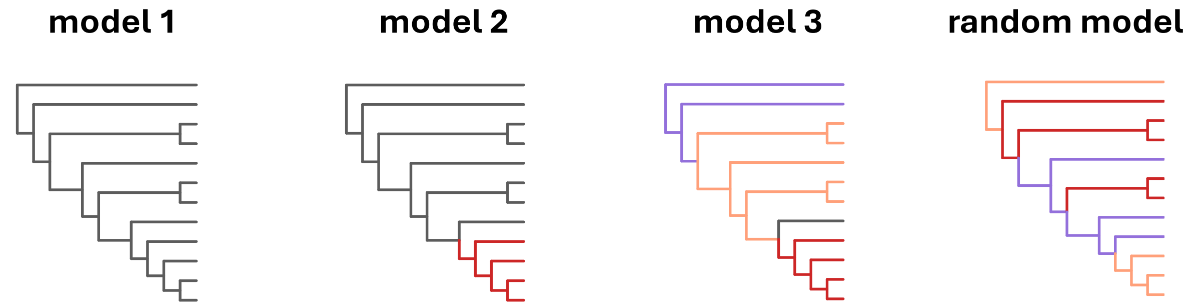


**Fig. S17. Four evolutionary models tested with CAGEE.**

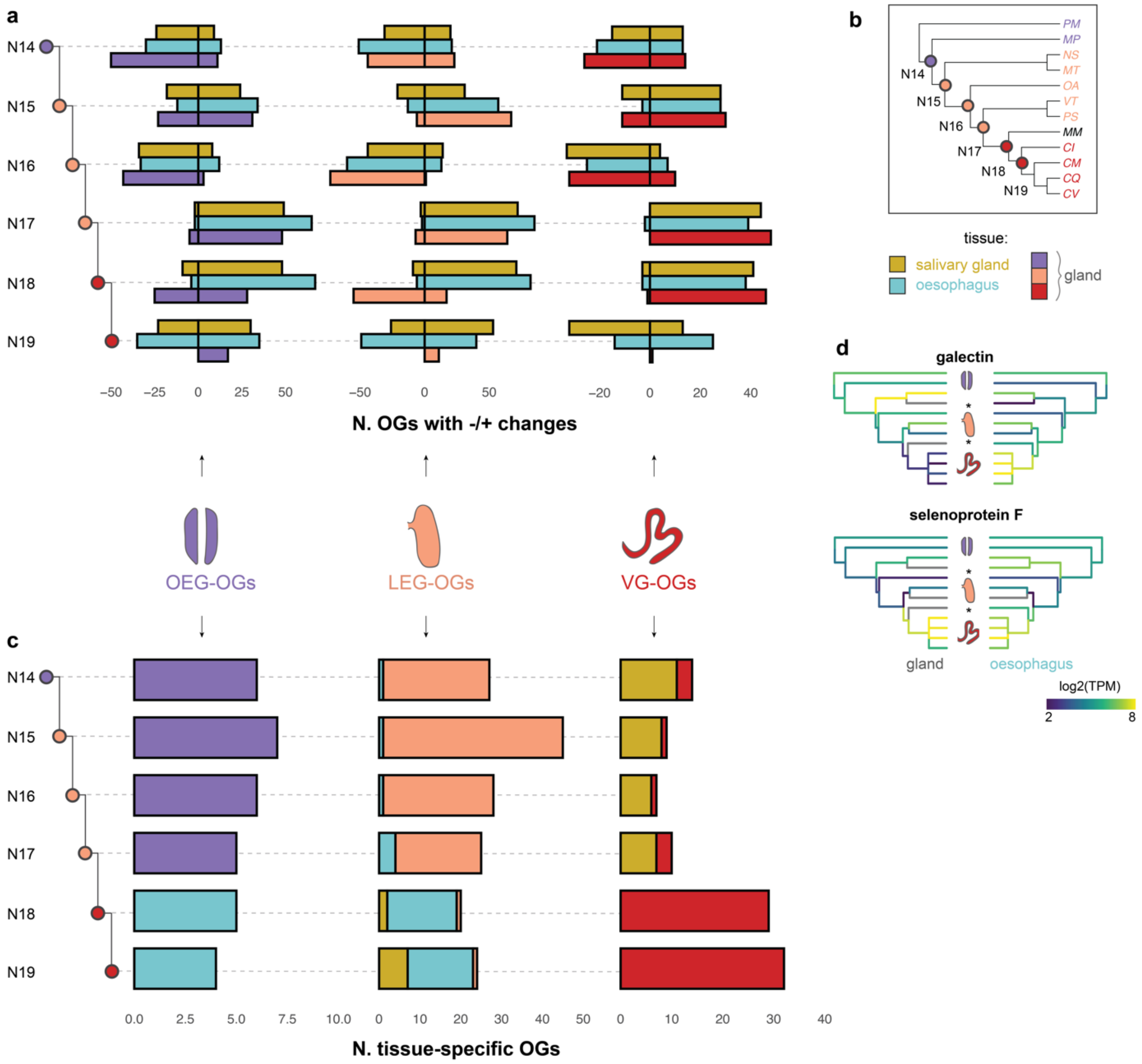


**Fig. S18. Gene expression dynamics across the phylogeny using the mean-based expression matrix.** a) Number of OEG-, LEG, and VG-specific OGs decreasing and increasing their expression levels in the ancestral salivary glands, oesophagus, or mid-oesophageal gland at each internal node of the gastropod phylogeny. The expression changes were calculated based on gene expression reconstruction at each node of the phylogeny. b) Species phylogeny with the number of the internal nodes. c) Number of OEG-, LEG-, and VG-specific OGs which were tissue-specific in the ancestral mid-oesophageal gland, salivary glands, and oesophagus at each node of the phylogeny. d) Ancestral reconstruction of gene expression of a galectin and selenoprotein F in the mid-oesophageal gland and oesophagus. The missing tissues are marked with a *. The orthogroup expression matrix used was based on the mean TPM across all the genes within an orthogroup.

**Table S1. Evolutionary rates for the four tested models.** For each organ is reported: the number of evolutionary rates estimated ($\sigma^2$), the likelihood, and the $\sigma^2$ estimates for each group. The orthogroup expression matrix used was based on the mean TPM across all the genes within an orthogroup.

| | model | N. of evolutionary rates (σ2) | likelihood (-ln L) | OEG-species σ2 | LEG-species σ2 | glandless-species σ2 | VG-species σ2 |
|---|---|---|---|---|---|---|---|
| gland | random | 3 | 17385.3 | | | | |
| 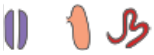 | 1 | 1 | 17891.7 | 0.97 | 0.97 | - | 0.97 |
| | 2 | 2 | 17097.3 | 0.61 | 0.61 | - | 1.57 |
| | **3** | **3** | **16725** | **0.28** | **0.74** | **-** | ***1.58*** |
| salivary glands | random | 3 | 17964.4 | | | | |
| | 1 | 1 | 18179.7 | 0.88 | 0.88 | 0.88 | 0.88 |
| | 2 | 2 | 16619 | 0.55 | 0.55 | 0.55 | 2.27 |
| | **3** | **3** | **16588.9** | **0.46** | **0.59** | **0.46** | ***2.26*** |
| oesophagus | random | 3 | 21770.8 | | | | |
| | 1 | 1 | 22154.2 | 0.99 | 0.99 | 0.99 | 0.99 |
| | 2 | 2 | 21425.4 | 0.70 | 0.70 | 0.70 | 1.57 |
| | **3** | **3** | **21277.4** | **0.46** | **0.80** | **0.46** | ***1.56*** |

**Table S2. Gastropod genomes included in the custom database used to annotate the *de novo* transcriptome assemblies.**

| species | genome ID | comments |
| --- | --- | --- |
| *Achatina fulica* | GCA_009760885.1 | http://gigadb.org/dataset/100647 |
| *Aplysia californica* | GCF_000002075.1 | |
| *Biomphalaria glabrata* | GCF_000457365.1 | |
| *Candidula unifasciata* | GCA_905116865.2 | |
| *Chrysomallon squamiferum* | GCA_012295275.1 | https://datadryad.org/stash/dataset/doi%253A 10.5061%252Fdryad.24053dn |
| *Lautoconus ventricosus* | GCA_018398815.1 | http://gigadb.org/dataset/100892 |
| *Elysia chlorotica* | GCA_003991915.1 | |
| *Gigantopelta aegis* | GCF_016097555.1 | |
| *Haliotis laevigata* | GCA_008038995.1 | https://abalonedb.org/genome-resources/data-downloads/ |
| *Lottia gigantea* | GCF_000327385.1 | |
| *Pomacea canaliculata* | GCF_003073045.1 | |

**Legend for Dataset S1.** List of the RNA-seq libraries created in this study and the relative sample information including species, specimen, tissue type, and number of reads.

**Legend for Dataset S2**. Overview of libraries and assembly statistics including the number of sequenced libraries per species, number of retained assembled transcripts after quality-filtering, percentage of completeness based on OMArk, percentage of contamination in the assemblies, and main contaminants.
